# Predicting metabolite response to dietary intervention using deep learning

**DOI:** 10.1101/2023.03.14.532589

**Authors:** Tong Wang, Hannah D. Holscher, Sergei Maslov, Frank B. Hu, Scott T. Weiss, Yang-Yu Liu

**Author notes:** **Correspondence**. Correspondence and requests for materials should be addressed to Y.-Y. L.

## Abstract

Due to highly personalized biological and lifestyle characteristics, different individuals may have different metabolite responses to specific foods and nutrients. In particular, the gut microbiota, a collection of trillions of microorganisms living in the gastrointestinal tract, is highly personalized and plays a key role in the metabolite responses to foods and nutrients. Accurately predicting metabolite responses to dietary interventions based on individuals’ gut microbial compositions holds great promise for precision nutrition. Existing prediction methods are typically limited to traditional machine learning models. Deep learning methods dedicated to such tasks are still lacking. Here we develop a method McMLP (**<u>M</u>**etabolite response predictor using **<u>c</u>**oupled **<u>M</u>**ulti**<u>l</u>**ayer **<u>P</u>**erceptrons) to fill in this gap. We provide clear evidence that McMLP outperforms existing methods on both synthetic data generated by the microbial consumer-resource model and real data obtained from six dietary intervention studies. Furthermore, we perform sensitivity analysis of McMLP to infer the tripartite food-microbe-metabolite interactions, which are then validated using the ground-truth (or literature evidence) for synthetic (or real) data, respectively. The presented tool has the potential to inform the design of microbiota-based personalized dietary strategies to achieve precision nutrition.

## Introduction

Precision nutrition aims to provide personalized dietary recommendations based on an individual’s unique biological and lifestyle characteristics such as genetics, gut microbiota, metabolomic profiles, and anthropometric data^1,2^. In addition to the design and implementation of large-scale clinical studies, one of the critical components for achieving precision nutrition is the development of predictive models that incorporate diverse individual data types to achieve an accurate prediction of metabolomic profiles following dietary changes^1–3^. However, existing models are limited to traditional machine learning methods such as Random Forest (RF)^4,5^ and Gradient-Boosting Regressor (GBR)^3^. Deep learning techniques have not been leveraged to predict metabolite responses for precision nutrition.

Among the biological characteristics relevant for precision nutrition, the gut microbiota is an important factor that explains a large fraction of individual metabolite responses among populations^4–7^. Indeed, the human gut microbiota produces many metabolites through the microbial metabolism of nondigested food components such as dietary fibers, which are prevalent in grains, vegetables and fruits^8^. Therefore, microbiota-derived metabolites are important mediators of host health^9–13^. For example, short-chain fatty acids (SCFAs) are metabolites produced by intestinal microbes through anaerobic fermentation of indigestible polysaccharides such as dietary fiber and resistant starch^9,10^. SCFA concentrations have been linked to regulation of immune cell function^14,15^, gut-brain communication^16^, and cardiovascular diseases^17,18^. Among the SCFAs, butyrate has been shown to be negatively correlated with pro-inflammatory cytokines^19,20^. Hence, a high level of butyrate from the gut microbiota is believed to be beneficial due to its anti-inflammatory effects^19–21^. Boosting the levels of health-beneficial metabolites by modulating the gut microbiota appears to be a promising approach to improve host health^22–24^.

One possible way to modulate the gut microbiota is through dietary interventions^6^. Gut microbial composition is affected by the diet^6,25–28^. As a result, microbiota-targeted dietary interventions have been proposed to modulate the gut microbiota to increase the production of metabolites beneficial to the host^29–31^. Recently, there has been a growing trend to exploit the tripartite relationship between food/nutrition, gut microbiota, and microbiota-derived metabolites to provide better dietary advice for each individual^3–5,28–32^. Indeed, accurate prediction of personalized metabolite responses to foods and nutrients based on the gut microbiota holds great promise for precision nutrition^33^.

Many dietary intervention studies have attempted to investigate the relationship between diet and microbial metabolism of the gut microbiota^27,28,32,34^. However, most of these studies only analyzed correlations between dietary treatments, microbes, or metabolites. A few studies have used different analytic approaches to predict postprandial responses of metabolite markers such as blood glucose^3,4^ and immune markers^4,34^. However, the personalized prediction of how important markers such as SCFAs and bile acids respond to long-term dietary interventions is under-investigated (Fig. 1).

**Figure 1:**
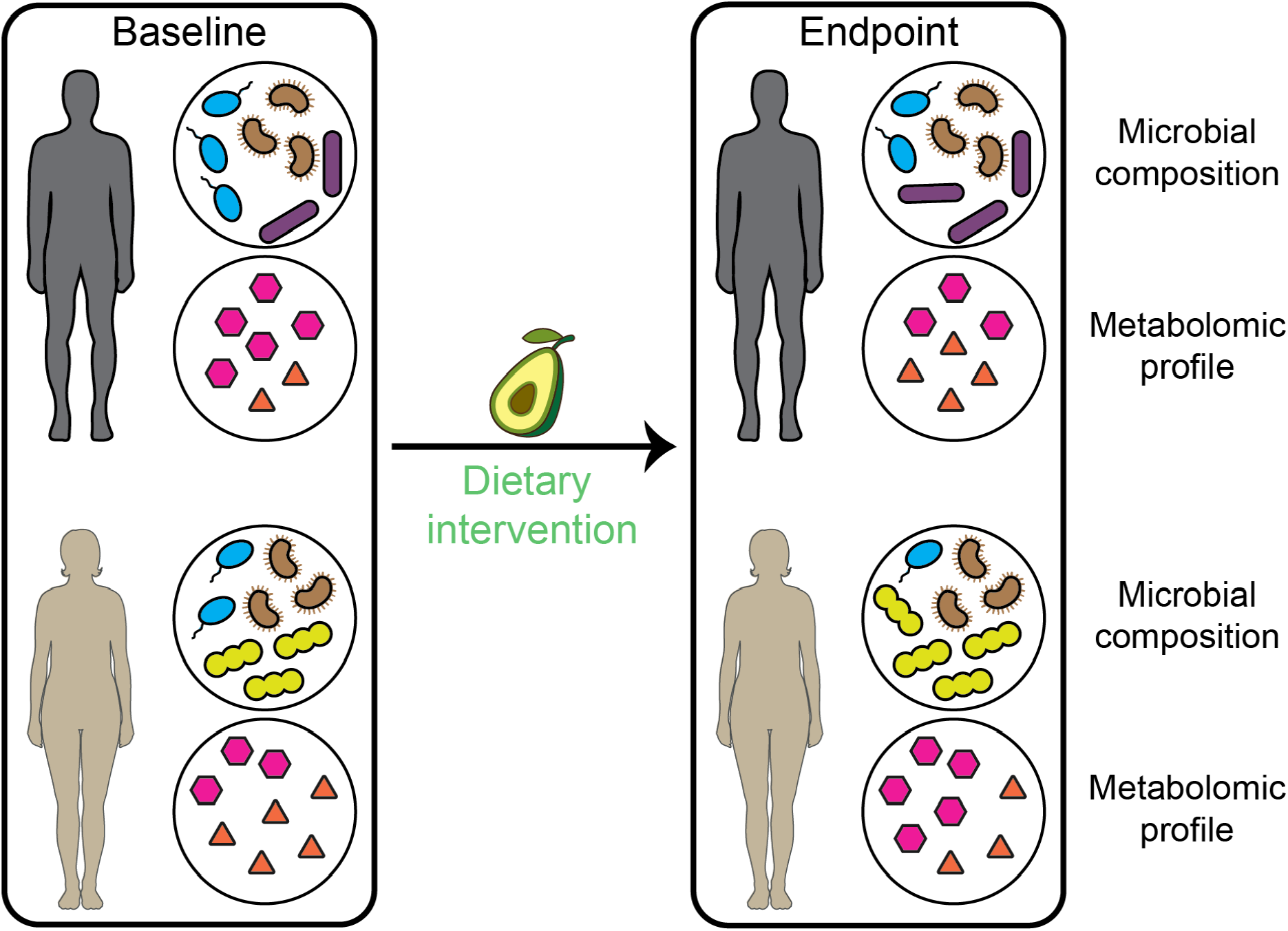
A typical dietary intervention study design. Before the dietary intervention, the baseline gut microbial compositions and metabolomic profiles (of either fecal samples or blood samples) are measured. During the dietary intervention, one or a few dietary resources are introduced (represented here by avocado) in addition to the baseline diet. The task we intend to solve is to predict personalized metabolite responses after dietary intervention based on the baseline gut microbial compositions, baseline metabolomic profiles, and the dietary intervention strategy.

Our aim is to predict the post-dietary intervention (or “endpoint”) metabolite concentrations in fecal or blood samples based on the pre-dietary intervention (or “baseline”) microbial composition, metabolome data, and the dietary intervention strategy. This is conceptually different from existing studies on the inference of metabolomic profiles from microbial compositions measured at the same time^35–38^. Herein, we leveraged data from randomized, controlled dietary intervention studies^27,28,34,39–41^ and developed a deep-learning method: **<u>M</u>**etabolite response predictor using **<u>c</u>**oupled **<u>M</u>**ulti**<u>l</u>**ayer **<u>P</u>**erceptrons (McMLP) to predict endpoint metabolite concentrations based on baseline microbial composition. We first generated synthetic data based on a microbial consumer-resource model that simulates the dietary intervention process and found that McMLP outperforms existing methods (RF and GBR), especially when the training sample size is small. We then applied all methods to real data from six dietary intervention studies^27,28,34,39–41^, finding that the predictive power of McMLP is higher than existing methods. Finally, based on the well-trained McMLP, we performed the sensitivity analysis to infer the tripartite food-microbe-metabolite relationship, supported by some literature evidence.

## Results

### Overview of McMLP

We hypothesized that in order to accurately predict post-dietary intervention metabolomic profiles, we first need to capture how the microbiome composition changes from the baseline to the endpoint. This is because metabolomic profiles reflect the microbial metabolism of a community^7,42^. To test our hypothesis, we proposed McMLP, which consists of two steps : (step-1) use the baseline microbiota and metabolome data (i.e., concentrations of targeted metabolites) and the dietary intervention strategy to predict the endpoint microbial composition; and (step-2) use the predicted endpoint microbial composition, the baseline metabolome data, and the dietary intervention strategy to predict the endpoint metabolomic profile (Fig. 2a; Supplementary Fig. 1a). For each step, we used a multilayer perceptron (MLP) with Rectified Linear Unit (ReLu) as the activation function to perform the prediction. We emphasize that, in principle, one can just use one MLP to directly predict endpoint metabolomic profiles based on baseline microbiota/metabolome data and the dietary intervention strategy (Supplementary Fig. 1b). Later, we confirmed that this one-step strategy has worse predictive power than our two-step strategy.

**Figure 2:**
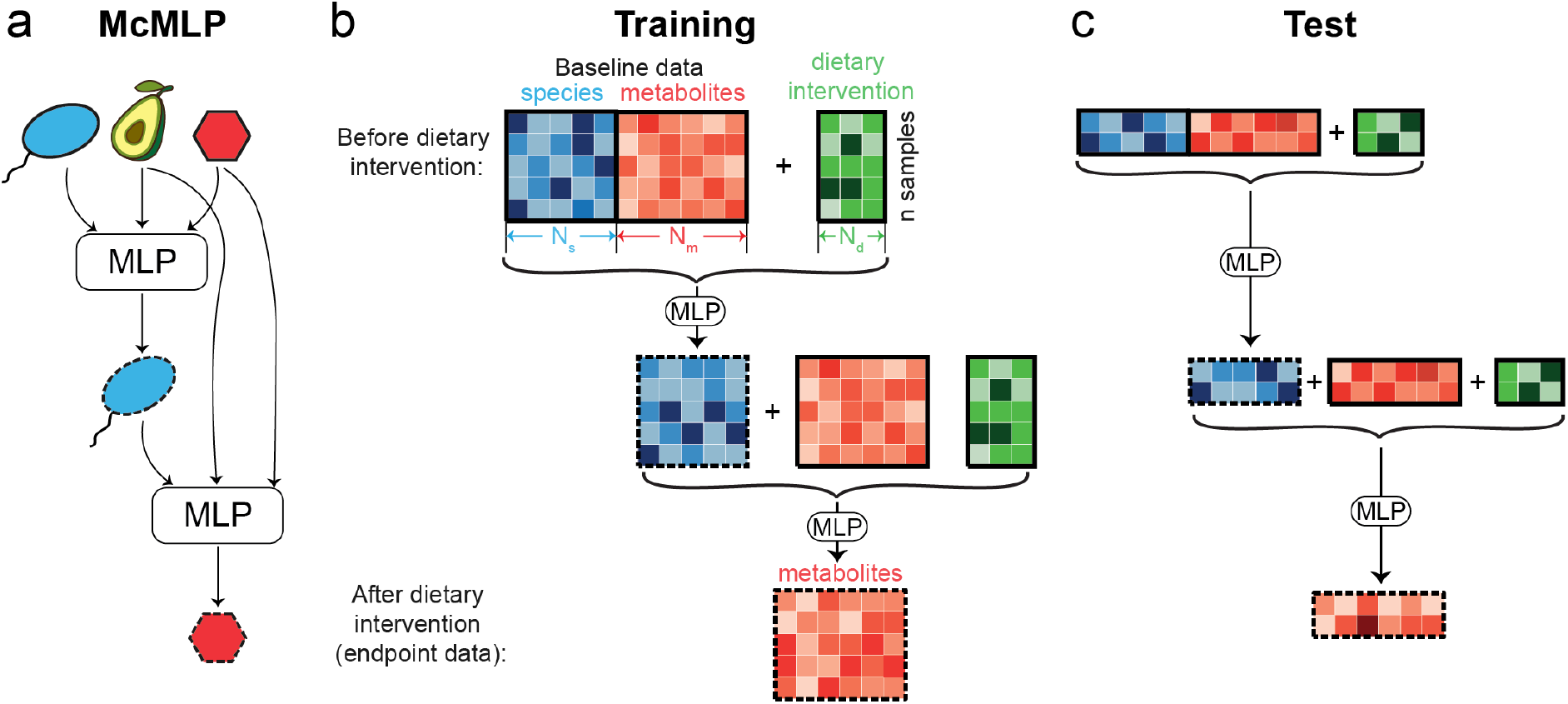
The workflow of McMLP (Metabolite response predictor using coupled multilayer perceptrons). We aim to predict endpoint metabolomic profiles (i.e., metabolomic profiles after the dietary interventions) based on the baseline microbial compositions (i.e., microbial compositions before the dietary intervention), dietary intervention strategy, and baseline metabolomic profiles. Here we used a hypothetical example with n=5 training samples and 2 samples in the test set. For each sample, we considered *N*_s_ microbial species, *N*_d_ dietary resources, and *N*_m_ metabolites. Across three panels, microbial species and their relative abundances are colored blue, dietary resources and their intervention doses are colored green, and metabolites and their concentrations are colored red. Icons associated with baseline/endpoint data are bounded by solid black/dashed lines respectively. **a**, The model architecture of McMLP. McMLP comprises two coupled MLPs (multilayer perceptrons). The first MLP at the top (step 1) predicts the endpoint microbial compositions based on the baseline data and the dietary intervention strategy. The predicted endpoint microbial compositions from the first MLP are then provided as input to the second MLP at the bottom (step 2). The second MLP combines the predicted endpoint microbial compositions, the dietary intervention strategy, and the baseline metabolomic profiles to finally predict the endpoint metabolomic profiles. The value of dietary intervention strategy is either binary to denote the presence/absence of each dietary resource or numeric to be proportional to the intervention dose. Details of both MLPs can be found in Supplementary Fig. 1 and Methods. **b**, McMLP takes two types of baseline data (baseline microbial compositions and baseline metabolomic profiles) and the dietary intervention strategy as input variables and is trained to predict corresponding endpoint metabolomic profiles. During training, the endpoint microbial composition is needed to train the first MLP. By contrast, the second MLP directly takes the predicted endpoint microbial composition instead of the actual endpoint microbial composition. **c**, The well-trained McMLP can generate predictions for metabolomic profiles for the test set. During testing, no endpoint microbial composition is needed because the second MLP directly takes the predicted endpoint microbial composition from the first MLP as the input.

From a practical standpoint, our goal is to predict an individual’s metabolite response (i.e., the change in concentrations of targeted metabolites) to a potential dietary intervention to facilitate precision nutrition. To achieve this goal, we feed the baseline microbiota and metabolome profiles of this individual and the potential dietary intervention strategy to a well-trained McMLP to predict the endpoint metabolome profile. Note that in this application (or test) stage, because the dietary intervention is a thought experiment, no real endpoint data is available. The first MLP in McMLP will predict the endpoint microbiota profile, which will be fed into the second MLP to predict the endpoint metabolome profile.

During the training stage of McMLP, we need to collect not only baseline microbiota and metabolome profiles of different individuals, but also perform dietary interventions to collect actual endpoint microbiota and metabolome profiles. We emphasize that the actual endpoint microbiota data will only be used to train the first MLP (Fig. 2b). It shall not be used to train the second MLP. This is because we need to keep the consistency between the training and application (or test) stages. After all, during the application stage, it is the predicted endpoint microbiome profile that will be fed into the second MLP, and the actual endpoint microbiome profile does not exist at all.

Instead of fine-tuning hyperparameters such as the number of layers *N*_1_ and the hidden layer dimension *N*_h_ for MLP, we overparameterized MLP by using a large and fixed number of layers *N*_1_ and hidden layer dimension *N*_h_ (*N*_1_ = 6 and *N*_h_ = 2048). The overparameterized machine learning methods, especially deep learning models, yield better performance due to their high capacity (i.e., more model parameters). In fact, the high-capacity models can be even simpler due to smoother function approximation and thus less likely to overfit^43^.

To illustrate the prediction task, we used a hypothetical example comprising *N*_s_(= 5) microbial species, *N*_d_(= 3) dietary resources being intervened, *N*_m_(= 6) metabolites, and 7 samples (Fig. 2b,c). We will use both the baseline data and the dietary intervention strategy as inputs for McMLP (Fig. 2a). We used the Centered Log-Ratio (CLR)-transformed microbial relative abundances as the microbial composition and log10 transformed metabolite concentrations as the metabolomic profile. We did not impose the constraint that the predicted relative abundances from the first MLP add up to one. The value of dietary intervention strategy is either binary to denote the presence/absence of each dietary resource or numeric to be proportional to the intervention dose. 5 samples are used as the training set (Fig. 2b) and the remaining 2 samples form the test set (Fig. 2c). To evaluate the regression performance, we employed three metrics based on the Spearman correlation coefficient (SCC) *ρ* between the predicted and true values of the concentration of one metabolite across all samples: (1)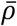: the mean SCC, (2) *f*_*ρ*>0.5_: the fraction of metabolites with *ρ* greater than 0.5, and (3)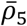: the mean SCC of the top-5 best-predicted metabolites.

### McMLP generates superior performance over existing methods on synthetic data

To validate the predictive power of McMLP, we applied it to synthetic data generated from the Microbial Consumer-Resource Model (MiCRM) which considers microbial interactions through both nutrient competition and metabolic cross-feeding^44^. We adapted MiCRM to simulate the dietary intervention. For simplicity, we considered 20 food resources, 20 microbes, and 20 metabolites in the modeling. Also, we assumed that food resources can only be consumed while metabolites can be either consumed or produced. Prior to the dietary intervention, one food resource (referred to as “food resource #1”) was not introduced, while the remaining 19 food resources were supplied. Dietary intervention was simulated by adding food resource #1 at a specific “dose” to microbial communities composed of surviving species before the dietary intervention and calculating the new ecological steady state. Here, the “dose” is defined as the ratio between the amount of the introduced food resource during the dietary intervention and the average amount of other food resources introduced before the dietary intervention. We split the synthetic data (with 250 samples) with 80/20 ratio fifty times to generate fifty train-test pairs that can be used to reflect the variation in predictive performance. Details on model simulation and synthetic data generation can be found in the Supplementary Information.

We compared the performance of McMLP with two classical methods (GBR: Gradient-Boosting Regressor^3^; RF: Random Forest^4,5^) in the prediction task defined in Fig. 2. For each method, we considered two sets of input variables: (1) without baseline metabolomic profiles (denoted as “w/o b” hereafter) and (2) with baseline metabolomic profiles (denoted as “w/ b” hereafter).

We first used the three metrics 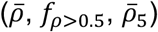 to benchmark the predictive performance of the different methods on synthetic data with 50 training samples and an intervention dose of 3. We found that McMLP generated the best performance (Figs. 3a1-a3), especially when baseline metabolomic profiles were included in the input. When we predict without baseline metabolomic profiles, McMLP is significantly better than RF and GBR (p-value < 0.05 for 5/6 comparison cases, Wilcoxon signed-rank test applied; McMLP yields the highest 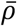 of 0.391 ± 0.008, the highest *f*_*ρ*>0.5_ of 0.19*7* ± 0.018, and the highest 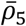of 0.536 ± 0.00*7*; the standard error is used to measure the variation in performance metrics across 50 train-test splits). Including baseline metabolomic profiles in the input significantly improves the performance of all methods, with McMLP still being the best (which yields the highest 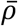of 0.595 ± 0.005, the highest *f*_*ρ*>0.5_ of 0.815 ± 0.014, and highest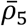 of 0.*7*15 ± 0.006). We also tried to introduce 5 food resources during the dietary intervention (instead of 1 previously; see Supplemental Information for details) and found that the performance of McMLP is still superior to other methods when the dietary intervention strategy is more complex (Supplementary Fig. 2).

**Figure 3:**
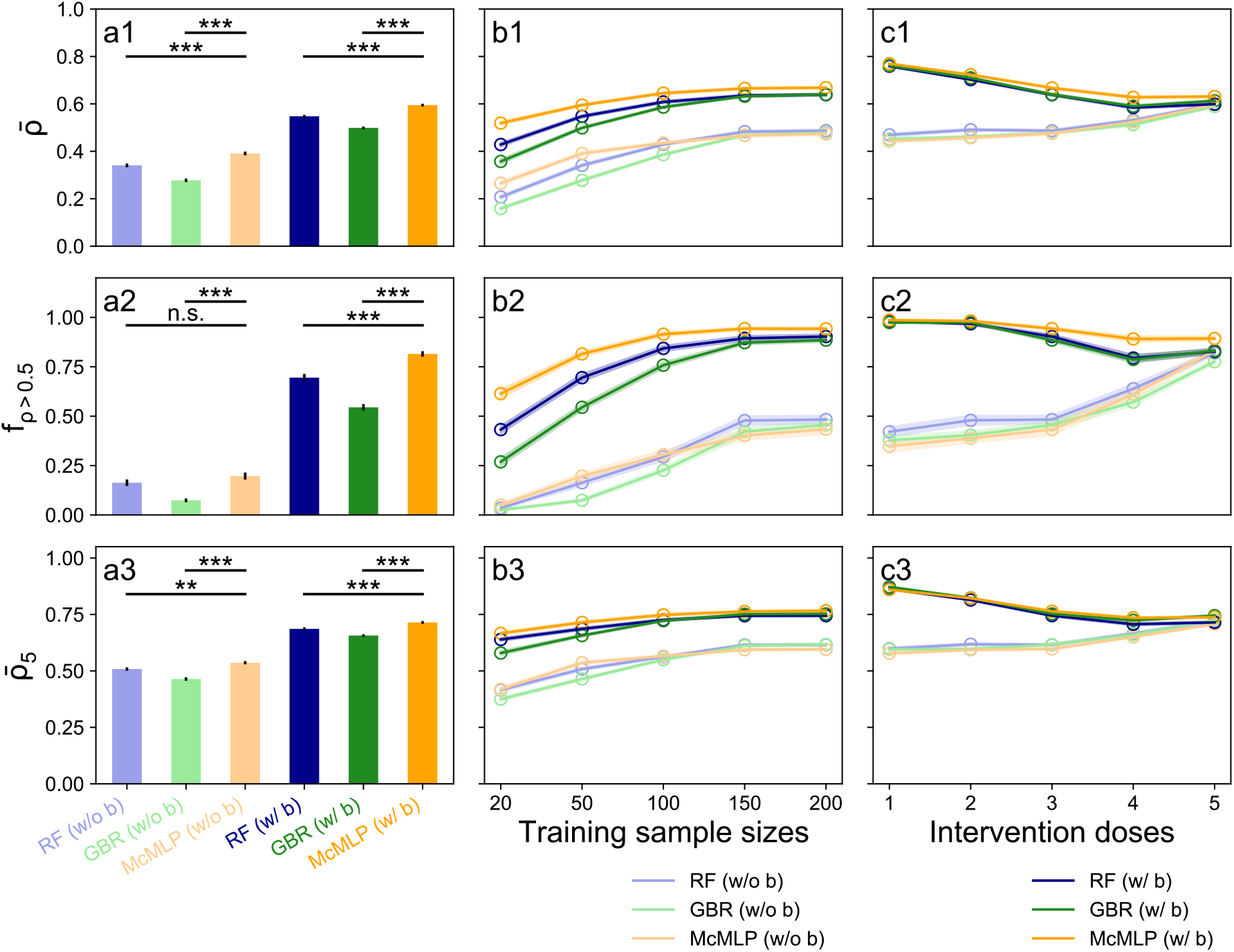
McMLP provides better predictive power than previously developed computational methods for predicting endpoint metabolomic profiles on synthetic data generated from microbial consumer-resource models. Three computational methods are compared: Random Forest (RF), Gradient Boosting Regressor (GBR), and McMLP. For each method, we either included (“w/ b” label) or did not include (“w/o b” label) baseline metabolomic profiles as input variables. Each method with a particular combination of input data is colored the same way in all panels. Standard errors are computed based on fifty random train-test splits and shown in all panels (as solid black vertical lines or transparent areas around their means). To compare different methods, we adopted three metrics: the mean Spearman Correlation Coefficient (SCC) 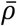, the fraction of metabolites with SCCs greater than 0.5 (denoted as *f*_*ρ*>0.5_), and the mean SCC of the top-5 predicted metabolites 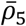. Error bars denote the standard error (n=50). **a1-a3**, For the synthetic data with an intervention dose of 3 and 50 training samples, McMLP provides the best performance for all three metrics regardless of whether the baseline metabolomic profiles are included or not. **b1-b3**, When the intervention dose is 3, the predictive performance of all methods gets better and closer to each other as the training sample size increases. Including baseline metabolomic profiles also helps to improve the prediction. **c1-c3**, When 200 training samples are used, the performance gap between including and not including baseline metabolomic profiles shrinks as the intervention dose increases. All statistical analyses were performed using the two-sided Wilcoxon signed-rank test. P values obtained from the test are divided into four groups: (1) *p* > 0.05 (n.s.), (2) 0.01 < *p* ≤ 0.05 (*), (3) 10^−3^ < *p* ≤ 0.01 (**), and (4) 10^−4^ < *p* ≤ 10^−3^ (***). Source data of raw data points and p values are provided as a Source Data file.

We further examined the effect of training sample size on model performance. While maintaining the same 50-sample test set used previously, we found that all performance metrics for all methods improved as the training sample size increased (Fig. 3b1-b3). More importantly, we found that the performance of McMLP is better than RF and GBR at small training sample sizes (20 or 50) and is close to RF and GBR at large training sample sizes (>50). This demonstrates the superior performance of McMLP with a limited number of samples, contrary to the traditional notion that deep learning methods tend to overfit at small sample sizes^45^.

We also examined the effect of intervention dose on model performance. By varying the concentration of the intervened food resource in MiCRM, we generated synthetic data with different intervention doses and subsequently trained all ML methods on them with 200 training samples. We found that the performance gap between methods using and not using baseline metabolomic profiles narrows as the intervention dose increases (Fig. 3c1-c3). We believe this is because a larger intervention dose significantly changes the endpoint metabolomic profile away from its baseline level, rendering the baseline metabolomic profile less useful.

Different from the above-mentioned benchmarking method where training data overlapped across train-test splits, we explored the impact of non-overlapping training data on our benchmarking results. To explore this, we created one independent synthetic dataset for each training and utilized the same, separate dataset as the test set (with 100 samples) for the performance evaluation across all repeats. Based on this new benchmarking protocol, we have benchmarked the performance of all algorithms and once again revealed the amazing predictive performance of McMLP (Supplementary Fig. 3).

### McMLP accurately predicts metabolite responses on real human gut microbiota data

After validating McMLP using synthetic data, we analyzed real data from six dietary intervention studies to see if its performance on real data was consistently better than existing methods. The first dataset we collected was from a study investigating how avocado consumption alters gut microbial compositions and concentrations of fecal metabolites such as SCFAs and bile acids^28^.

In this study all participants were divided into two groups based on the food components of the meals provided: (1) avocado group: 175 g (men) or 140 g (women) of avocado was provided as part of a meal once a day for 12 weeks and (2) control group: no avocado was included in their control meal^28^. Baseline (i.e., before the dietary intervention) and endpoint (i.e., during week 12 of the intervention) microbial compositions and concentrations of SCFAs and bile acids were quantified. The dataset is unique due to its relatively large sample size (66 for both avocado and control groups)^28^ compared to other dietary intervention studies^27,32,34^.

Because the amount of avocado consumed by participants in the avocado group was very similar and participants in the control group barely consumed avocado, for simplicity, we encoded the participant’s dietary intervention in McMLP and other methods as a binary variable in the input (green icons/symbols representing diets in Fig. 2) whose value equals 1 or 0 if the participant is in the avocado or control group, respectively. Note that in this study the concentrations of fecal SCFAs and bile acids were obtained from two separate targeted metabolomic assays. Hence, we separated the concentration prediction of SCFAs and bile acids to compare the predictability of the two metabolite classes. We found that for the concentration prediction of both SCFAs and bile acids, McMLP with the baseline metabolomic profiles consistently produces the best performance (Fig. 4a1-a3, b1-b3). Interestingly, the inclusion of baseline metabolomic profiles in the input of McMLP helps more with the prediction of bile acid concentrations than with the prediction of SCFA concentrations (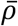increases from 0.182 to 0.346 for bile acids when metabolomic profiles are included; 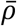increases from 0.260 to 0.262 for SCFAs when metabolomic profiles are included). A potential explanation is that the correlation of SCFA concentrations between baseline and endpoint samples is weaker than that of bile acids (Supplementary Fig. 4).

**Figure 4:**
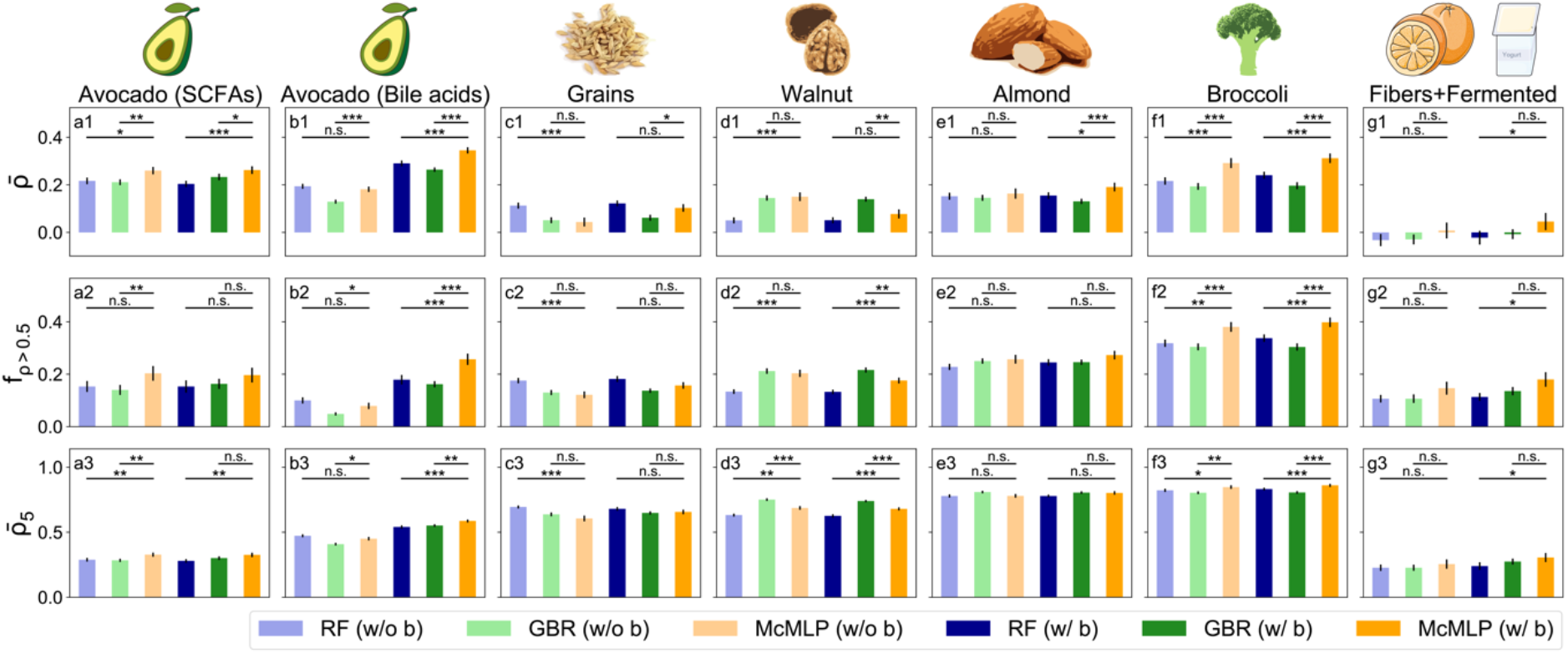
McMLP is superior to previous methods in terms of predicting endpoint metabolomic profiles on real data from six dietary intervention studies. Three computational methods are compared: Random Forest (RF), Gradient Boosting Regressor (GBR), and McMLP. For each method, we either included (“w/ b” label) or did not include (“w/o b” label) baseline metabolomic profiles as input variables. Each method with a particular combination of input data is colored the same in all panels. Standard errors are computed based on fifty random train-test splits and shown in all panels (solid black vertical lines). To compare different methods, we adopted three metrics: the mean Spearman Correlation Coefficient (SCC) 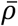, the fraction of metabolites with SCCs greater than 0.5 (denoted as *f*_*ρ*>0.5_, and the mean SCC of the top-5 predicted metabolites 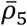. Error bars denote the standard error (n=50). **a1-a3**, Comparison of the performance in predicting SCFAs on the data from the avocado intervention study^28^. **b1-b3**, Comparison of performance in predicting bile acids on the data from the avocado intervention study^28^. **c1-c3**, Comparison of predictive performance on the data from the grain intervention study^39^. **d1-d3**, Comparison of predictive performance on the data from the walnut intervention study^27^. **e1-e3**, Comparison of predictive performance on the data from the almond intervention study^40^. **f1-f3**, Comparison of predictive performance on the data from the broccoli intervention study^41^. **g1-g3**, Comparison of predictive performance on the data from the high-fiber food or fermented food intervention study^34^. All statistical analyses were performed using the two-sided Wilcoxon signed-rank test. P values obtained from the test are divided into four groups: (1) *p* > 0.05 (n.s.), (2) 0.01 < *p* ≤ 0.05 (*), (3) 10^−3^ < *p* ≤ 0.01 (**), and (4) 10^−4^ < *p* ≤ 10^−3^ (***). Source data of raw data points and p values are provided as a Source Data file.

We checked the predictive performance of the one-step strategy that uses the same number of layers and nodes as one step in McMLP (*N*_1_ = 6 and *N*_h_ = 2048 in Supplementary Fig. 1b), finding that it is not as good as that of McMLP (Supplementary Fig. 5). It is worth noting that augmenting the one-step approach with additional data types through the two-step McMLP does not automatically guarantee enhanced predictive performance. The utility of the additional data hinges on its relevance and the model’s capacity to utilize it efficiently. Despite these potential uncertainties, we believe the enhanced performance of McMLP could be attributed to its two-step approach. This method allows for an initial capture of the endpoint microbial composition, presumably better associated with the endpoint metabolite concentrations. This may also explain why McMLP outperforms RF^4,5^ and GBR^3^, which employ a one-step approach and do not leverage the endpoint microbial compositions during method training. We also compared McMLP with the state-of-art method of predicting metabolomic profiles from microbial compositions measured at the same time --- mNODE^38^, finding that it has a worse performance than McMLP (Supplementary Fig. 6). The worse performance of mNODE is likely due to the fact that it is not dedicated to predicting metabolomic profiles at different time points. More technical reasons can be found in the Supplementary Information.

We extended the method comparison to five additional datasets from independent dietary studies investigating how microbiota compositions and fecal metabolite concentrations were influenced by adding grains^39^, walnuts^27^, almonds^40^, broccoli^41^, and high-fiber or fermented foods^34^ (the number of fecal microbes and metabolites as well as the types of metabolites are summarized in Table 1; see Methods section for details of the studies). Each participant’s dietary intake was similarly encoded as either a binary variable or a vector whose value is proportional to the consumed amount of the added dietary component, depending on the complexity of the dietary intervention. Further details of the data processing and model architecture setup can be found in the Supplementary Information. As shown in Fig. 4, McMLP consistently produces the best performance across all datasets (p-value < 0.05 for 47/84 comparison cases, Wilcoxon signed-rank test applied). The relatively poor performance of all methods on the data from the study that investigated fibers and fermented foods^34^ is likely due to the fact that a variety of foods within the fiber and fermented foods categories were consumed by the participants at will, while other studies were complete feeding trials^34^.

**Table 1:** Summary of key features of dietary intervention studies used in our method comparison. ASVs: Amplicon Sequence Variants.

| Dietary Intervention Studies | # of participants | # of intervention periods or groups | # of ASVs | # of fecal metabolites | Types of fecal metabolites |
| --- | --- | --- | --- | --- | --- |
| **Avocado**<sup>28</sup> | 132 | 2 | 278 | 27 | SCFAs, BCFAs, and bile acids |
| **Grains**<sup>39</sup> | 68 | 3 | 650 | 43 | Amino acids, carboxylic acids, lipids, bile acids, and monosaccharides |
| **Walnut**<sup>27</sup> | 18 | 2 | 419 | 41 | Amino acids, carboxylic acids, lipids, bile acids, and monosaccharides |
| **Almond**<sup>40</sup> | 18 | 5 | 714 | 43 | Amino acids, carboxylic acids, lipids, and bile acids |
| **Broccoli**<sup>41</sup> | 18 | 2 | 855 | 35 | Amino acids, carboxylic acids, lipids, and bile acids |
| **Fibers or fermented foods**<sup>34</sup> | 32 | 2 | 503 | 9 | SCFAs and BCFAs |

We noticed that the predictive performance of McMLP on real data is worse than that in synthetic data. We believe the observed discrepancy in predictive performance between the synthetic and real data may be due to the influence of human host, such as host metabolism^46^ and health status^47^. While 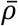appears to be low (∼0.2 to 0.4), the top-5 best-predicted metabolites for each dataset have great predictability, likely due to their strong association with the gut microbiome (Supplementary Fig. 7). We also compared the predictive performance of McMLP with that of a simple MLP with one hidden layer with everything else the same as in McMLP, finding that McMLP generates better performance (Supplementary Fig. 8).

We also explored whether incorporating covariates in the metadata can help further improve the predictive performance of McMLP. We only obtained the covariates for the avocado intervention study. For the avocado dataset, we have three covariates: gender, BMI, and age. We included these three covariates as additional variables in McMLP, finding that the incorporation of covariates significantly improves the predictive performance for most cases (Fig. 5). We also analyzed the permutation feature importance of the three covariates by shuffling the values of a covariate in the input and then measuring the reduction in the average Spearman Correlation Coefficients 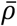. We found that all three covariates are important, except that gender is slightly less important than age when predicting the SCFAs (Supplementary Fig. 9).

**Figure 5:**
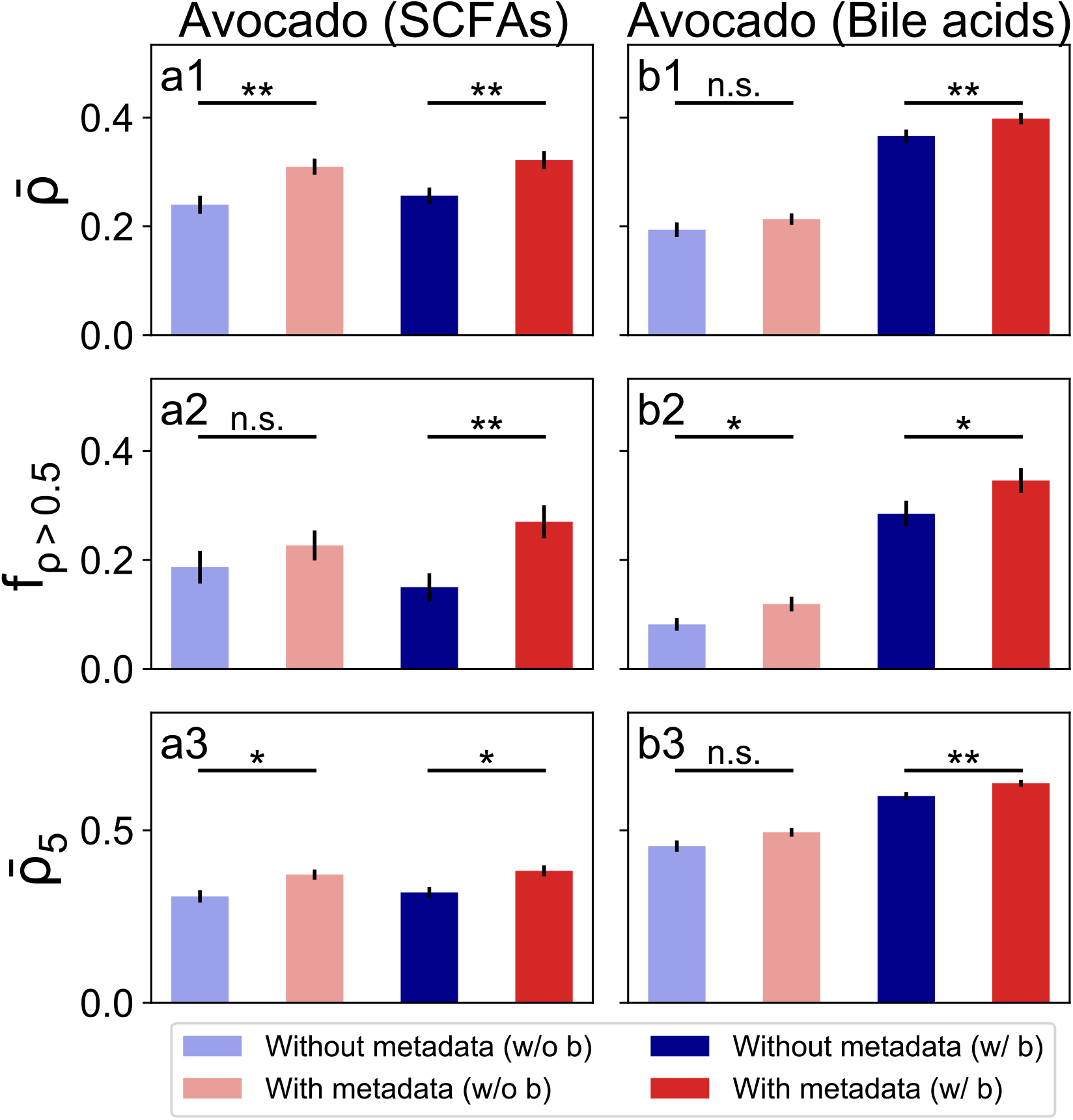
Including the covariates in metadata (age, BMI, and gender) in the input of McMLP improves it in terms of predicting endpoint metabolomic profiles on real data from the avocado intervention study. All results are derived from McMLP. We either included (“w/ b” label) or did not include (“w/o b” label) baseline metabolomic profiles as input variables. Each method with a particular combination of input data is colored the same in all panels. Standard errors are computed based on fifty random train-test splits and shown in all panels (solid black vertical lines). To compare different methods, we adopted three metrics: the mean Spearman Correlation Coefficient (SCC) 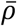, the fraction of metabolites with SCCs greater than 0.5 (denoted as *f*_*ρ*>0.5_), and the mean SCC of the top-5 predicted metabolites 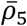. Error bars denote the standard error (n=50). **a1-a3**, Comparison of the performance in predicting SCFAs on the data from the avocado intervention study^28^. **b1-b3**, Comparison of performance in predicting bile acids on the data from the avocado intervention study^28^. All statistical analyses were performed using the two-sided Wilcoxon signed-rank test. P values obtained from the test are divided into four groups: (1) *p* > 0.05 (n.s.), (2) 0.01 < *p* ≤ 0.05 (*), (3) 10^−3^ < *p* ≤ 0.01 (**), and (4) 10^−4^ < *p* ≤ 10^−3^ (***). Source data of raw data points and p values are provided as a Source Data file.

We wonder if the predictive performance of McMLP can be enhanced if we use the functional profiles generated from the whole-metagenome shotgun (WMS) sequencing instead of the microbial compositions derived from the 16S rRNA gene sequencing. To test this, we leveraged the available WMS sequencing data for a subset of samples in the avocado study. In the end, only 45 individuals have paired baseline-endpoint data. Their functional profiles are represented by 375 pathway features (see Methods section for details). For the 45 paired baseline-endpoint data, we compared the predictive performance among three different input data types: (1) microbial compositions, (2) functional profiles, and (3) combining both microbial compositions and functional profiles. The performance comparison of the three different input data types yields no significant difference (Supplementary Figs. 10 and 11).

For the avocado dataset, we also grouped the ASV (Amplicon Sequence Variants) compositions from the 16S rRNA gene sequencing and the species-level microbial compositions from the WMS sequencing to the genus level. When analyzing the 16S sequencing data, predictions using the ASV-level compositions are generally more accurate than those using the genus-level compositions (Supplementary Fig. 12). For SCFAs, the predictive performances based on two types of compositions are comparable. Regarding the WMS data, we observed that predictions using the species-level compositions are slightly better than those using the genus-level compositions (Supplementary Fig. 13).

### Inferring the tripartite food-microbe-metabolite relationship

It has been previously shown that an individual’s metabolite response depends on her/his gut microbial composition^7,42,48^. If we want to introduce a new dietary resource to boost the concentration of a health-beneficial metabolite mediated by gut microbes, we need to target “key” microbial species that meet two criteria: (1) the species can consume one or more dietary components in the introduced food resource; (2) the species can increase the metabolite concentration. If either criterion is not met, it is difficult to boost the metabolite concentration via this dietary intervention. Specifically, we identify these “key” species that satisfy both criteria by revealing the food-microbe consumption and microbe-metabolite production patterns, which can be summarized in a tripartite food-microbe-metabolite graph (Supplementary Fig. 14). To achieve this, we performed the sensitivity analysis of McMLP. In particular, we interpreted a potential relationship between an input variable *x* and an output variable *y* by perturbing *x* by a small amount (denoted as Δ*x*) and then measuring the response of *y* (denoted as Δ*y*). Following the notion of sensitivity in engineering sciences, we defined sensitivity 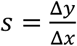 and used its sign (positive/negative) to reflect whether *y* changes in the same/opposite direction as *x*. More technical details of this calculation can be found in the Methods section or in our previous study^38^. We calculated sensitivities for step-1 (and step-2) in McMLP to infer potential food-microbe consumption (and microbe-metabolite production) interactions, respectively (Fig. 6a). Specifically, in step-1, we perturbed the amount of food resource *α* and measured the change in the relative abundance of species *i*. The sensitivity of species *i* to food resource *α* is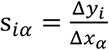 and its sign can be used to reflect the interaction between species *i* and food resource *α*. s_*iα*>_ 0, indicates that species *i* can consume some nutrient components of food resource *α*. Similarly, for step-2, we define the sensitivity of metabolite *β* to species *i* as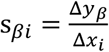. The positive sensitivity, s_*βi*_ >0, reveals potential production of the metabolite *β* by species *i*.

**Figure 6:**
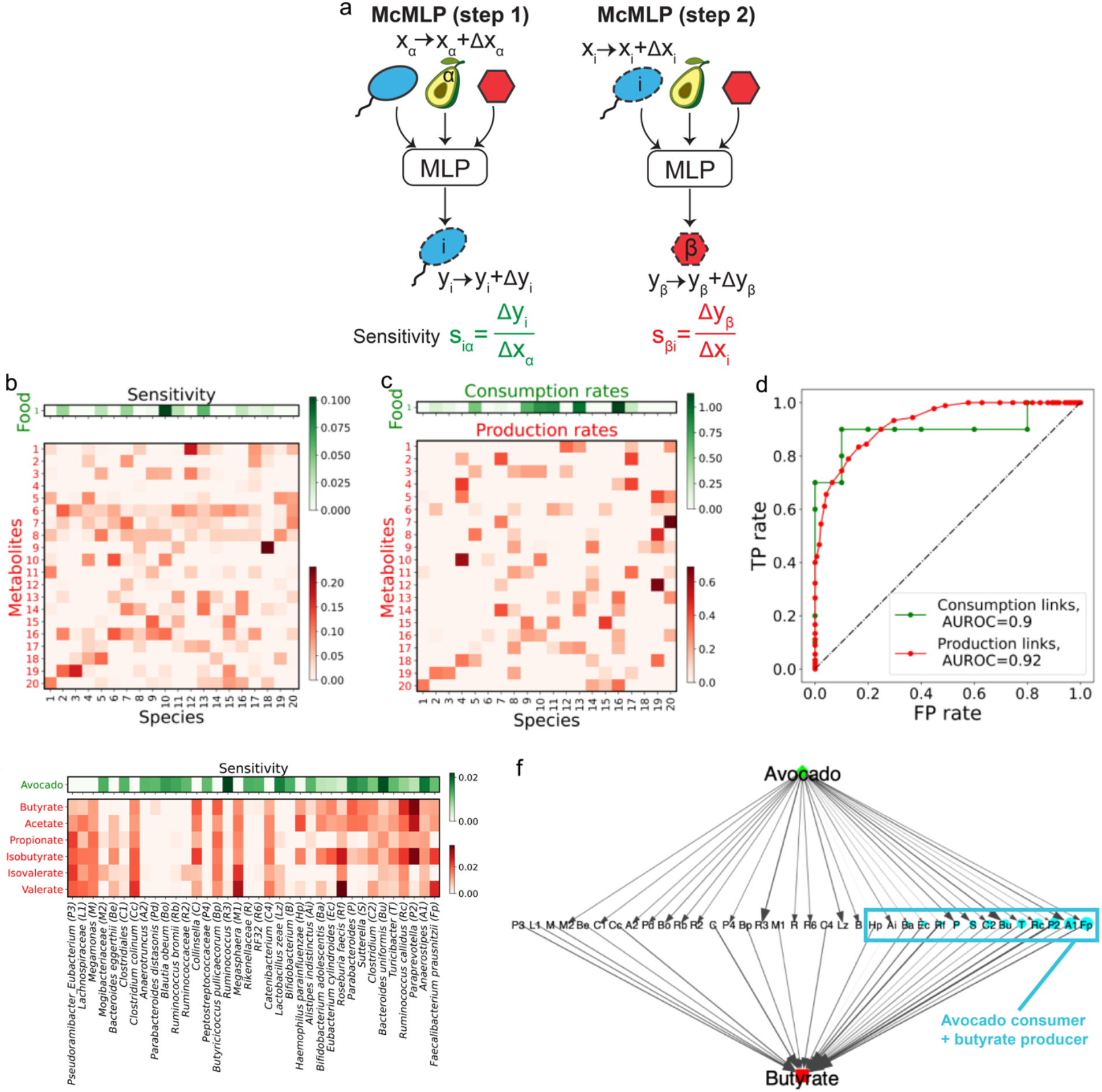
Applying sensitivity analysis of McMLP accurately infers food-microbe consumption interactions and microbe-metabolite production interactions in both synthetic and real data. **a**, The sensitivity of the relative abundance of species *i* to the supplied dietary resource *α* is denoted as s_*iα*_. It is defined as the ratio between the change in the relative abundance of species *i* (Δ_*i*._) and a small perturbation in the supplied dietary resource *α* (Δ*x*_*α*_). Similarly, the sensitivity of the concentration of metabolite *β* to the relative abundance of species *i* is denoted as s_*βi*._. It is defined as the ratio between the change in the concentration of metabolite *β* (Δ*y*_*β*_) and the perturbation in the relative abundance of species *i* (Δ*x*_*i*_). **b**, The sensitivity values for food-microbe consumption interactions (colored in green) and microbe-metabolite production interactions (colored in red) in the synthetic data. **c**, The ground-truth food-microbe consumption rates (colored in green) and microbe-metabolite production rates (colored in red) in the synthetic data. **d**, The Area Under the Receiver Operating Characteristic (AUROC) curve based on True Positive (TP) rates and False Positive (FP) rates which are obtained by using different sensitivity thresholds to classify interactions. **e**, The sensitivity values for avocado-microbe consumption interactions (colored in green) and microbe-metabolite production interactions (colored in red) for the real data from the avocado intervention study. **f**, The avocado-microbe-butyrate tripartite graph constructed based on the sensitivity values of avocado-microbe consumption interactions and microbe-butyrate production interactions for the real data from the avocado intervention study. The edge width and edge arrow sizes are proportional to the absolute values of the sensitivities. All microbes in the middle layer are arranged from left to right in the increasing order of the incoming edge width multiplied by the outgoing edge width. Source data are provided as a Source Data file.

We first evaluated our sensitivity method on the synthetic data for which we know the ground truth of food-microbe consumption and microbe-metabolite production interactions. We found that the inferred sensitivity values for all food-microbe and microbe-metabolite pairs (Fig. 6b) have a zero-nonzero pattern very similar to the ground-truth consumption and production rates assigned in MiCRM (Fig. 6c). We chose zero as the sensitivity threshold and kept only positive values for food-microbe pairs (green cells in Fig. 6b&c) and for microbe-metabolite pairs (red cells in Fig. 6b&c) to explore consumption and production interactions respectively. To statistically verify the agreement between ground-truth interactions and inferred interactions based on sensitivity values, we computed the AUROC (Area Under the Receiver Operating Characteristic curve) based on the overlap between true and predicted interactions when the classification threshold is varied. More specifically, for each classification threshold *s*_thres_, we predicted the consumption of food resource *α* by species *i* (or production of metabolite *α* by species *i*) to be true only if *s*_*αi*_ > *s*_thres_ (or *s*_*αi*_> *s*_thres_). We achieved excellent performance in inferring either food-microbe consumption interactions (green line and dots with AUROC=0.9 in Fig. 6d) or microbe-metabolite production interactions (red line and dots with AUROC=0.92 in Fig. 6d).

We then performed the same inference on real data from the avocado study^28^. The results are shown in Fig. 6e. (Inference results of other studies are provided in the Supplementary Tables.) Our results shown in Fig. 6e are in agreement with prior biological knowledge that *Faecalibacterium prausnitzii* is a stronger producer of butyrate^49^ than *Ruminococcus callidus*, and *R. calidus* is a stronger producer of acetate than *F. prausnitzii*^50,51^.

The inference results also enable us to construct the tripartite food-microbe-metabolite graph. For the sake of simplicity, here we visualize the avocado-microbe-butyrate subgraph (Fig. 6f). Note that increased butyrate levels have been shown to be beneficial to host health by enhancing immune status^19–21^. For the avocado-microbe-butyrate subgraph, we focused on the top-20 avacado-microbe consumption and top-20 microbe-butyrate production interactions ranked by their absolute sensitivity values. Only nodes and links associated with these interactions were shown in this subgraph. Widths of individual edges in this figure are proportional to the absolute values of the corresponding sensitivities and node sizes for microbes are proportional to the products of edge widths connecting this microbe to avocado at the top and butyrate at the bottom of this subgraph. We ordered microbial nodes in the middle layer in the increasing order of node sizes from left to right (Fig. 6f). This organization helps us identify the key species that serve as both strong consumers of avocado and strong producers of butyrate. *F. prausnitzii* emerged as the most important key species for butyrate production in response to avocado intervention. Our results are consistent with previous studies^49^. For example, *F. prausnitzii* levels have been previously shown to be elevated when avocado is supplied by diet^52^. In a separate study, *F. prausnitzii* has also been shown to produce butyrate as a metabolic byproduct^49^.

## Discussion

A highly accurate computational method for predicting metabolite responses based on baseline data and a potential dietary intervention strategy is a prerequisite for precision nutrition. In this paper, we developed a deep learning method, McMLP, which predicts metabolomic profiles after a dietary intervention better than existing methods. We first validated the superior performance of McMLP using synthetic data generated by a microbial consumer-resource model and investigated the influence of diet intervention doses and training sample sizes. We then demonstrated that McMLP produced the most accurate predictions across six different dietary intervention studies^27,28,34,39–41^. We proceeded with a biological interpretation of McMLP results using sensitivity analysis to infer the tripartite food-microbe-metabolite relationship, finding that the inferred relationship was quite accurate in synthetic data. Finally, we demonstrated that our sensitivity analysis applied to real data revealed key species whose metabolic capabilities were consistent with prior biological knowledge.

Currently available dietary intervention studies have many limitations for use in machine learning. First, the sample size (or number of participants) of these studies is typically small, on the order of dozens^27,32,34,41^. The relatively small sample size fundamentally limits the performance of any predictive model. While the cross-validation that we employed is a widely used method for assessing model robustness and preventing overfitting, its reliability is contingent upon the sample size. It has been well-documented that performance estimates derived from cross-validation can carry a significant degree of uncertainty when applied to small datasets such as dietary intervention with walnuts, almonds, and broccoli^53^. This uncertainty is attributed to the increased variability in training and validation splits, which can result in overestimated or underestimated model performance. This problem may be mitigated in ongoing large-scale research cohorts with many participants. One such cohort is the All of Us Research Program, which is attempting to build a diverse health database of more than one million people across the U.S. and then use the data to learn how human biology, lifestyle, and environment affect health. As part of this observational cohort, the recently announced Nutrition for Precision Health Study will recruit 10,000 participants to conduct precision dietary interventions^54^. Second, only a handful of dietary components have ever been explored in dedicated diet-microbiota studies. As a result, the computational approaches can only predict metabolite responses for the limited set of dietary components used in these studies. However, to realize the promise of precision nutrition to provide accurate personalized dietary recommendations, we need a predictive model that can accurately predict metabolite responses for a wide range of dietary components. Last, other baseline variables unavailable to us here (e.g., meal composition, age, sex, demographics, and anthropometric data) might help to improve the predictive performance. If such data are available, they can be incorporated into McMLP as extra input variables.

Consistent with most dietary intervention studies in the literature, the data available for this study were primarily the 16S rRNA gene sequencing data. We acknowledge that 16S rRNA gene sequencing may restrict our taxonomic resolution to the genus level for certain taxa. Yet, a unique aspect of this work is that the 16S results were able to be further explored and validated using WMS data that were available for a subset of the avocado study. Our comparison of the predictive performance of McMLP between using the microbial compositions and the functional profiles in the input demonstrates the effectiveness of using the microbial compositions (at the ASV level) derived from the 16S rRNA gene sequencing data. Indeed, McMLP still yields highly promising results for identifying important interactions supported by works of literature: (1) *Faecalibacterium prausnitzii* is a stronger producer of butyrate and (2) *Ruminococcus callidus* is a stronger producer of acetate. Both *Faecalibacterium prausnitzii* and *Ruminococcus callidus* are identified from the 16S data. These results showcase the power of performing the sensitivity analysis on well-trained McMLP. Looking ahead, we believe this lack of data will only be solved by the emergence of more datasets from dietary intervention studies with paired metabolome and WMS sequencing data. Hopefully, the emergence of new datasets in the future will open new opportunities for applying and refining our method. We recognize the inherent complexity of microbe-metabolite interactions and acknowledge that our approach primarily focuses on very simplified pairwise consumption/production interactions between microbes and metabolites. In reality, these interactions can extend beyond pairwise interactions. For example, the serum level of enterolactone, one metabolite produced by some gut microbiota upon consumption of dietary fibers, is influenced by the dietary fat intake^56^. Additionally, McMLP currently relies on a limited selection of metabolites obtained from targeted metabolomics. In the future, as dietary intervention studies incorporate more comprehensive lists of metabolites, we anticipate that the prediction power of McMLP will be further improved.

Our McMLP architecture is quite generic --- its input variables and their dimensions can be easily adapted to fit more complex datasets. For example, if a particular dietary intervention study documents an extensive list of dietary components, McMLP can be modified to include an input node for each dietary component to reflect the amount and frequency of its consumption. Similarly, the predicted output variables of McMLP need not be limited to metabolomic profiles measured in fecal samples. It can be generalized to predict other variables such as immune biomarkers or metabolite concentrations from blood samples. Moreover, McMLP can be interpreted through sensitivity analysis, revealing numerous interactions supported by existing literature evidence. If McMLP can successfully predict other data types, it might be feasible to infer other types of interactions.

Unlike other machine learning methods that typically require hyperparameter tuning to achieve the best performance for each dataset with a different set of hyperparameters, McMLP consistently outperformed existing machine learning methods across six real datasets even without hyperparameter tuning. We speculate that McMLP exploited the recently observed “double-descent” behavior for the risk curve^43^, which suggests that an overparametrized deep-learning model (i.e., one with an extremely large number of model parameters) can generate better and more consistent performance than models with less capacity and more carefully tuned hyperparameters. To reach this overparameterized regime, we used a large and fixed number of layers *N*_1_ = 6 and a large hidden layer dimension *N*_h_ = 2048, exceeding both the number of microbial species and the number of metabolites. One benefit of using such a model free of hyperparameter tuning is the shorter training time. Since the typical 5-fold cross-validation used to select the best set of hyperparameters is the most time-consuming part of a typical deep learning workflow, McMLP saves a significant amount of time required for hyperparameter tuning and thus has a shorter training time (∼ 5 minutes for each run of McMLP on the avocado intervention study^28^).

## Methods

### Ethical Compliance Statement

In this study, we used publicly available datasets from previous studies, whose study procedures were administered in accordance with the Declaration of Helsinki and were approved by the University of Illinois Institutional Review Board.

### Datasets

The datasets utilized herein were generated as part of work on bacterial^56^ and metabolite^57^ biomarkers of food intake, which provided anonymized microbial and metabolomic data. The dataset related to the fibers or fermented foods intervention study is available for download in the supplemental material of the original publication^34^. The main characteristics of the dietary intervention studies used above are summarized in Table 1. Across all studies, fecal or blood samples were collected before and after each dietary intervention period. Gut microbiota composition was determined by the 16S rRNA gene sequencing and metabolomic profiles of either fecal samples or blood serum samples were determined by tandem liquid chromatography-mass spectrometry (LC-MS/MS) and gas chromatography-mass spectrometry (GC-MS) metabolomics. For all machine learning tasks, the same fifty random 80/20 train-test splits were used to ensure a fair comparison of methods. Further details are described below:

Avocado intervention study. This dataset was reported by a dietary intervention study that investigated how avocado consumption altered the relative abundance of gut bacteria and concentrations of microbial metabolites in 132 adults with overweight or obesity^28^. All participants were assigned to the avocado treatment or no-avocado control group (66 each for arm). They consumed isocaloric meals with or without avocado (175 g, men; 140 g, women) once daily for 12 weeks. For fecal samples collected before and after the dietary intervention, 278 ASVs (Amplicon Sequence Variants) were determined by the 16S rRNA gene sequencing and profiles of 6 SCFAs and 21 bile acids were generated by LC-MS/MS metabolomics. Out of 132 individuals, 45 individuals’ fecal samples (both collected before and after the dietary intervention) underwent whole-metagenome shotgun (WMS) sequencing. To obtain the taxonomic profiles, DIAMOND (double index alignment of next-generation sequencing data) v2.0.11.149 was used in conjunction with the National Center for Biotechnology Information (NCBI) non-redundant (nr) protein reference database (June 2021) to align translated DNA query sequences. DIAMOND was set to “sensitive” mode, targeting alignments with >40% identity with an e-value of 0.00001^58^. Each sample’s sequences from the merged and cleaned 165 FASTQ file were aligned against the NCBI-nr database, producing a corresponding output DIAMOND alignment archive file. DIAMOND was set to “sensitive” mode, targeting 167 alignments with >40% identity with an e-value of 0.00001^58^. To generate the functional profiles, MEGAN (MEtaGenome ANalyzer) v6.12.2 Ultimate Edition was then used to perform functional analysis of the sequence alignments against the KEGG gene database^58^. For each sample, the sequence alignments produced by DIAMOND in the previous step were matched to a KEGG ortholog (KO) accession, producing a MEGAN file containing the total count of each KO across each sample^58^. NCBI taxonomy counts were also exported from MEGAN in a similar fashion^58^. Eventually, 859 microbial species-level microbial taxa were identified and 14,109 KOs were identified in the functional profiles. Since the number of KOs (14,109) is too large to be included in our machine-learning algorithms, all KOs are grouped into pathways. Eventually, there are 375 pathway features in the functional profiles.

Grains intervention study. This dietary intervention study investigated how grain barley and oat consumption affects gut bacteria relative abundances and concentrations of microbial metabolites in 68 healthy adults^39^. All participants were randomly assigned to receive one of three treatments: (1) a control diet containing 0.8 daily servings of whole grain/1800 kcal, (2) a diet containing 4.4 daily servings of whole grain barley/1800 kcal or (3) a diet containing 4.4 daily servings of whole grain oats/1800 kcal. Fecal samples were collected before and after the dietary intervention.

Walnut intervention study. This dietary intervention study investigated how walnut consumption affects the gut microbiota and metabolite concentrations in 18 healthy adults^27^. All participants completed two 3-week treatment/intervention periods separated by a 1-week washout period. Fecal samples were collected before and after the dietary intervention period.

Almond intervention study. This dietary intervention study was conducted in 18 healthy adults^40^. All participants completed four 3-week treatment periods and one control period separated by a 1-week washout period. Fecal samples were collected before and after the dietary intervention period.

Broccoli intervention study. In this study, 18 healthy adults completed two 18-day treatment periods separated by a 24-day washout period^41^. Fecal samples were collected before and after the dietary intervention period.

Fibers or fermented foods intervention study. This dietary intervention study was designed to investigate how consumption of plant-based foods rich in dietary fibers or fermented foods alters gut bacteria and their associated metabolites in 36 healthy adults^34^. All participants were divided into the high-fiber or the high-fermented-foods arm (18 each for arm). The entire dietary intervention lasted 17 weeks. Their fecal or blood serum samples were collected before and after the dietary intervention period. Gut microbiota composition in fecal samples was determined by the 16S rRNA gene sequencing and metabolomic profiles of serum samples were generated by the LC-MS metabolomics.

### McMLP

McMLP consists of two coupled MLPs: (step-1) in the first step (using the MLP at the top in Supplementary Fig. 1a), we predict endpoint microbial compositions based on baseline microbial compositions, baseline metabolomic profiles, and dietary intervention strategy; (step-2) in the second step (using the MLP at the bottom in Supplementary Fig. 1a), we take the predicted endpoint microbial compositions from the first MLP, baseline metabolomic profiles, and dietary intervention strategy to predict endpoint metabolomic profiles.

- Data processing: The CLR (Centered Log-Ratio) transformation is applied to microbial relative abundances and the log10 transformation is applied to metabolite concentrations.
- Model detail: Each MLP model (for either the top or the bottom MLP in Supplementary Fig. 1) has 6 hidden layers in the middle, sandwiched by input and output variables. Each hidden layer has a fixed hidden layer dimension of 2048.
- Training method: The Adam optimizer^59^ is used for the gradient descent. Specifically, each dataset is split into the train-test set with 80/20. Then the training set is further split into the train-validation set with 80/20. Training stops when (1) the Mean Squared Error (MSE) averaged over all metabolites on the training set is less than 0.1 and (2) the mean SCC (Spearman Correlation Coefficient) of annotated metabolites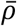on the validation set starts to decrease within the last 20 epochs.
- Activation function: ReLU (Rectified Linear Unit).

### Inference of food-microbe and microbe-metabolite interactions via sensitivity

The two MLP models in the well-trained McMLP can be interpreted separately. We first interpret the first MLP (step 1) in McMLP for food-microbe consumption interactions by the amount of food resource *α* (Δ*x*_*α*_) and then measure the change in the relative abundance of species *i* (Δ*y*_*i*_). Mathematically, for the sample *m* in the training set, we set the new value of this variable as zero. As a result, the perturbation amount for this variable in sample *m* is 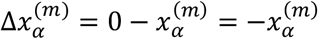where 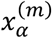is the unperturbed value. We can measure the change in the relative abundance of species *i* for sample 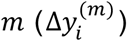 and define the sensitivity of species *i* to food resource *α* for sample *m* as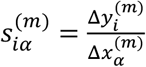. Finally, we can average sensitivity values across samples to obtain the average sensitivity of species *i* to food resource 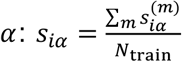 where *N*_train_ is the number of training samples. Similarly, for the second MLP (step-2) in McMLP, we can define 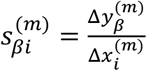and to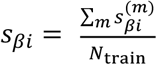 infer microbe-metabolite interactions by perturbing the relative abundance of species *i* (Δ*x*_*i*_) and then measuring the change in concentration of metabolite *β* (Δ*y*_*β*_).

### Statistics

To calculate correlations throughout the study, we used Spearman’s correlation coefficient. Wherever P-values were used we calculated the associated null distributions were computed from scratch. The non-parametric Wilcoxon signed-rank test is employed to evaluate differences in predictive performance among the algorithms. All statistical tests were performed using standard numerical and scientific computing libraries in the Python programming language (version 3.7.1) and Jupyter Notebook (version 6.1).

## Data Availability

Instructions for downloading 16S rRNA gene sequencing and metabolomics data analyzed in this work can be found in the literature exploring bacterial biomarkers of food intake^56^, metabolite biomarkers^57^, and the effects of fibers or fermented foods on immune markers^34^. To facilitate the data downloading, the URLs to those datasets are also provided in our McMLP GitHub repository (https://github.com/wt1005203/McMLP)^60^. The shotgun metagenomic sequencing data and covariates of the avocado intervention study, which belong to a separate research project and have not yet been published, will be made accessible by the PI of the avocado intervention study Dr. Hannah Holscher upon reasonable request. Source data are provided with this paper.

## Code Availability

The code of McMLP was deposited to the same McMLP GitHub repository.

## Acknowledgements

Y.-Y.L. is supported by grants R01AI141529, R01HD093761, RF1AG067744, UH3OD023268, U19AI095219 and U01HL089856 from the National Institutes of Health, USA; a pilot grant from the Biology of Trauma Initiative of Broad Institute, USA; and the Office of the Assistant Secretary of Defense for Health Affairs, through the Traumatic Brain Injury and Psychological Health Research Program (Focused Program Award) under award no. (W81XWH-22-S-TBIPH2), endorsed by the Department of Defense, USA. H.D.H has received grant funding from the Hass Avocado Board.

## Author Contributions

Y.-Y.L. conceived the project. T.W. and Y.-Y.L. designed the project.

T.W. performed all the numerical calculations and data analysis. All authors analyzed the results.

T.W. and Y.Y.L. wrote the manuscript. All authors edited and approved the manuscript.

## References

1. Hurtado-Lorenzo, A., Honig, G. & Heller, C. Precision Nutrition Initiative: Toward Personalized Diet Recommendations for Patients With Inflammatory Bowel Diseases. Crohn’s & Colitis 360 2, otaa087 (2020).

2. Bush, C. L., Blumberg, J. B., El-Sohemy, A., Minich, D. M., Ordovás, J. M., Reed, D. G., & Behm, V. A. Y. Toward the Definition of Personalized Nutrition: A Proposal by The American Nutrition Association. Journal of the American College of Nutrition 39, 5–15 (2020).

3. Zeevi, D., Korem, T., Zmora, N., Israeli, D., Rothschild, D., Weinberger, A., et al. Personalized Nutrition by Prediction of Glycemic Responses. Cell 163, 1079–1094 (2015).

4. Berry, S. E., Valdes, A. M., Drew, D. A., Asnicar, F., Mazidi, M., Wolf, J., et al. Human postprandial responses to food and potential for precision nutrition. Nat Med 26, 964–973 (2020).

5. Asnicar, F., Berry, S. E., Valdes, A. M., Nguyen, L. H., Piccinno, G., Drew, D. A., et al. Microbiome connections with host metabolism and habitual diet from 1,098 deeply phenotyped individuals. Nat Med 27, 321–332 (2021).

6. Kolodziejczyk, A. A., Zheng, D. & Elinav, E. Diet–microbiota interactions and personalized nutrition. Nat Rev Microbiol 17, 742–753 (2019).

7. Chen, L., Zhernakova, D. V., Kurilshikov, A., Andreu-Sánchez, S., Wang, D., Augustijn, H. E., et al. Influence of the microbiome, diet and genetics on inter-individual variation in the human plasma metabolome. Nat Med 1–11 (2022) doi:10.1038/s41591-022-02014-8.

8. Holscher, H. D. Dietary fiber and prebiotics and the gastrointestinal microbiota. Gut Microbes 8, 172–184 (2017).

9. Donia, M. S. & Fischbach, M. A. Small molecules from the human microbiota. Science 349, 1254766 (2015).

10. Koh, A., De Vadder, F., Kovatcheva-Datchary, P. & Bäckhed, F. From Dietary Fiber to Host Physiology: Short-Chain Fatty Acids as Key Bacterial Metabolites. Cell 165, 1332–1345 (2016).

11. Koppel, N., Maini Rekdal, V. & Balskus, E. P. Chemical transformation of xenobiotics by the human gut microbiota. Science 356, eaag2770 (2017).

12. Myhrstad, M. C. W., Tunsjø, H., Charnock, C. & Telle-Hansen, V. H. Dietary Fiber, Gut Microbiota, and Metabolic Regulation—Current Status in Human Randomized Trials. Nutrients 12, 859 (2020).

13. Cong, J., Zhou, P. & Zhang, R. Intestinal Microbiota-Derived Short Chain Fatty Acids in Host Health and Disease. Nutrients 14, 1977 (2022).

14. Corrêa-Oliveira, R., Fachi, J. L., Vieira, A., Sato, F. T. & Vinolo, M. A. R. Regulation of immune cell function by short-chain fatty acids. Clin Transl Immunology 5, e73 (2016).

15. Parada Venegas, D., De la Fuente, M.K., Landskron, G., González, M. J., Quera, R., Dijkstra, G., et al. Short Chain Fatty Acids (SCFAs)-Mediated Gut Epithelial and Immune Regulation and Its Relevance for Inflammatory Bowel Diseases. Frontiers in Immunology 10, (2019).

16. Dalile, B., Van Oudenhove, L., Vervliet, B. & Verbeke, K. The role of short-chain fatty acids in microbiota–gut–brain communication. Nat Rev Gastroenterol Hepatol 16, 461–478 (2019).

17. Richards, L. B., Li, M., van Esch, B. C. A. M., Garssen, J. & Folkerts, G. The effects of short-chain fatty acids on the cardiovascular system. PharmaNutrition 4, 68–111 (2016).

18. Ohira, H., Tsutsui, W. & Fujioka, Y. Are Short Chain Fatty Acids in Gut Microbiota Defensive Players for Inflammation and Atherosclerosis? Journal of Atherosclerosis and Thrombosis 24, 660–672 (2017).

19. Segain, J. P., De La Blétiere, D. R., Bourreille, A., Leray, V., Gervois, N., Rosales, C., et al. Butyrate inhibits inflammatory responses through NFκB inhibition: implications for Crohn’s disease. Gut 47, 397–403 (2000).

20. Säemann, M. D., Böhmig, G. A., Österreicher, C. H., Burtscher, H., Parolini, O., Diakos, C., et al. Anti-inflammatory effects of sodium butyrate on human monocytes: potent inhibition of IL-12 and up-regulation of IL-10 production. The FASEB Journal 14, 2380–2382 (2000).

21. He, J., Chu, Y., Li, J., Meng, Q., Liu, Y., Jin, J., et al. Intestinal butyrate-metabolizing species contribute to autoantibody production and bone erosion in rheumatoid arthritis. Science Advances 8, eabm1511 (2022).

22. Hughes, R. L., Marco, M. L., Hughes, J. P., Keim, N. L. & Kable, M. E. The Role of the Gut Microbiome in Predicting Response to Diet and the Development of Precision Nutrition Models—Part I: Overview of Current Methods. Advances in Nutrition 10, 953–978 (2019).

23. Hughes, R. L., Kable, M. E., Marco, M. & Keim, N. L. The Role of the Gut Microbiome in Predicting Response to Diet and the Development of Precision Nutrition Models. Part II: Results. Advances in Nutrition 10, 979–998 (2019).

24. Senior, M. Precision nutrition to boost cancer treatments. Nature Biotechnology 40, 1422– 1424 (2022).

25. David, L. A., Maurice, C. F., Carmody, R. N., Gootenberg, D. B., Button, J. E., Wolfe, B. E., et al. Diet rapidly and reproducibly alters the human gut microbiome. Nature 505, 559–563 (2014).

26. Rajoka, M. S. R., Shi, J., Mehwish, H. M., Zhu, J., Li, Q., Shao, D., et al. Interaction between diet composition and gut microbiota and its impact on gastrointestinal tract health. Food Science and Human Wellness 6, 121–130 (2017).

27. Holscher, H. D., Guetterman, H. M., Swanson, K. S., An, R., Matthan, N. R., Lichtenstein, A. H., et al. Walnut Consumption Alters the Gastrointestinal Microbiota, Microbially Derived Secondary Bile Acids, and Health Markers in Healthy Adults: A Randomized Controlled Trial. The Journal of Nutrition 148, 861–867 (2018).

28. Thompson, S. V., Bailey, M. A., Taylor, A. M., Kaczmarek, J. L., Mysonhimer, A. R., Edwards, C. G., et al. Avocado Consumption Alters Gastrointestinal Bacteria Abundance and Microbial Metabolite Concentrations among Adults with Overweight or Obesity: A Randomized Controlled Trial. The Journal of Nutrition 151, 753–762 (2021).

29. Suez, J. & Elinav, E. The path towards microbiome-based metabolite treatment. Nat Microbiol 2, 1–5 (2017).

30. Wong, A. C. & Levy, M. New Approaches to Microbiome-Based Therapies. mSystems 4, e00122–19 (2019).

31. Gulliver, E. L., Young, R. B., Chonwerawong, M., D’Adamo, G. L., Thomason, T., Widdop, J. T., et al. The future of microbiome-based therapeutics. Alimentary Pharmacology & Therapeutics 56, 192–208 (2022).

32. Deehan, E. C., Yang, C., Perez-Muñoz, M. E., Nguyen, N. K., Cheng, C. C., Triador, L., et al. Precision Microbiome Modulation with Discrete Dietary Fiber Structures Directs Short-Chain Fatty Acid Production. Cell Host & Microbe 27, 389-404.e6 (2020).

33. Wang, D. D. & Hu, F. B. Precision nutrition for prevention and management of type 2 diabetes. The Lancet Diabetes & Endocrinology 6, 416–426 (2018).

34. Wastyk, H. C., Fragiadakis, G. K., Perelman, D., Dahan, D., Merrill, B. D., Feiqiao, B. Y., et al. Gut-microbiota-targeted diets modulate human immune status. Cell 184, 4137–4153 (2021).

35. Mallick, H., Franzosa, E. A., Mclver, L. J., Banerjee, S., Sirota-Madi, A., Kostic, A. D., et al. Predictive metabolomic profiling of microbial communities using amplicon or metagenomic sequences. Nature Communications 10, 1–11 (2019).

36. Reiman, D., Layden, B. T. & Dai, Y. MiMeNet: Exploring microbiome-metabolome relationships using neural networks. PLOS Computational Biology 17, e1009021 (2021).

37. Le, V., Quinn, T. P., Tran, T. & Venkatesh, S. Deep in the Bowel: Highly Interpretable Neural Encoder-Decoder Networks Predict Gut Metabolites from Gut Microbiome. BMC Genomics 21, 256 (2020).

38. Wang, T., Wang, X. W., Lee-Sarwar, K. A., Litonjua, A. A., Weiss, S. T., Sun, Y., et al. Predicting metabolomic profiles from microbial composition through neural ordinary differential equations. Nat Mach Intell 5, 284–293 (2023).

39. Thompson, S. V., Swanson, K. S., Novotny, J. A., Baer, D. J. & Holscher, H. D. Gastrointestinal Microbial Changes Following Whole Grain Barley and Oat Consumption in Healthy Men and Women. The FASEB Journal 30, 406.1-406.1 (2016).

40. Holscher, H. D., Taylor, A. M., Swanson, K. S., Novotny, J. A. & Baer, D. J. Almond Consumption and Processing Affects the Composition of the Gastrointestinal Microbiota of Healthy Adult Men and Women: A Randomized Controlled Trial. Nutrients 10, 126 (2018).

41. Kaczmarek, J. L., Liu, X., Charron, C. S., Novotny, J. A., Jeffery, E. H., Seifried, H. E., et al. Broccoli consumption affects the human gastrointestinal microbiota. The Journal of Nutritional Biochemistry 63, 27–34 (2019).

42. Zierer, J., Jackson, M. A., Kastenmüller, G., Mangino, M., Long, T., Telenti, A., et al. The fecal metabolome as a functional readout of the gut microbiome. Nat Genet 50, 790–795 (2018).

43. Belkin, M., Hsu, D., Ma, S. & Mandal, S. Reconciling modern machine-learning practice and the classical bias–variance trade-off. PNAS 116, 15849–15854 (2019).

44. Marsland III, R., Cui, W., Goldford, J., Sanchez, A., Korolev, K., & Mehta, P. Available energy fluxes drive a transition in the diversity, stability, and functional structure of microbial communities. PLoS Comput Biol 15, e1006793 (2019).

45. James, G., Witten, D., Hastie, T., & Tibshirani, R. An Introduction to Statistical Learning, New York: springer 112, 18 (2013).

46. Visconti, A., Le Roy, C. I., Rosa, F., Rossi, N., Martin, T. C., Mohney, R. P., et al. Interplay between the human gut microbiome and host metabolism. Nat Commun 10, 4505 (2019).

47. Duboc, H., Rajca, S., Rainteau, D., Benarous, D., Maubert, M. A., Quervain, E., et al. Connecting dysbiosis, bile-acid dysmetabolism and gut inflammation in inflammatory bowel diseases. Gut 62, 531–539 (2013).

48. Dekkers, K. F., Sayols-Baixeras, S., Baldanzi, G., Nowak, C., Hammar, U., Nguyen, D., et al. An online atlas of human plasma metabolite signatures of gut microbiome composition. Nat Commun 13, 5370 (2022).

49. Zhou, L., Zhang, M., Wang, Y., Dorfman, R. G., Liu, H., Yu, T., et al. Faecalibacterium prausnitzii Produces Butyrate to Maintain Th17/Treg Balance and to Ameliorate Colorectal Colitis by Inhibiting Histone Deacetylase 1. Inflammatory Bowel Diseases 24, 1926–1940 (2018).

50. Cremon, C., Guglielmetti, S., Gargari, G., Taverniti, V., Castellazzi, A. M., Valsecchi, C., et al. Effect of Lactobacillus paracasei CNCM I-1572 on symptoms, gut microbiota, short chain fatty acids, and immune activation in patients with irritable bowel syndrome: A pilot randomized clinical trial. United European Gastroenterology Journal 6, 604–613 (2018).

51. Gaffney, J., Embree, J., Gilmore, S. & Embree, M. 2022. Ruminococcus bovis sp. nov., a novel species of amylolytic Ruminococcus isolated from the rumen of a dairy cow. International Journal of Systematic and Evolutionary Microbiology 71, 004924.

52. Cires, M. J., Navarrete, P., Pastene, E., Carrasco-Pozo, C., Valenzuela, R., Medina, D. A., et al. Protective Effect of an Avocado Peel Polyphenolic Extract Rich in Proanthocyanidins on the Alterations of Colonic Homeostasis Induced by a High-Protein Diet. J. Agric. Food Chem. 67, 11616–11626 (2019).

53. Isaksson, A., Wallman, M., Göransson, H. & Gustafsson, M. G. Cross-validation and bootstrapping are unreliable in small sample classification. Pattern Recognition Letters 29, 1960– 1965 (2008).

54. National Institutes of Health. NIH awards $170 million for precision nutrition study. National Institutes of Health (NIH) https://www.nih.gov/news-events/news-releases/nih-awards-170-million-precision-nutrition-study (2022).

55. Stumpf, K., Pietinen, P., Puska, P. & Adlercreutz, H. Changes in Serum Enterolactone, Genistein, and Daidzein in a Dietary Intervention Study in Finland1. Cancer Epidemiology, Biomarkers & Prevention 9, 1369–1372 (2000).

56. Shinn, L. M., Li, Y., Mansharamani, A., Auvil, L. S., Welge, M. E., Bushell, C., et al. Fecal Bacteria as Biomarkers for Predicting Food Intake in Healthy Adults. The Journal of Nutrition 151, 423–433 (2021).

57. Shinn, L. M., Mansharamani, A., Baer, D. J., Novotny, J. A., Charron, C. S., Khan, N. A., et al. Fecal Metabolites as Biomarkers for Predicting Food Intake by Healthy Adults. The Journal of Nutrition nxac195 (2022) doi:10.1093/jn/nxac195.

58. Shinn, L. M., Mansharamani, A., Baer, D. J., Novotny, J. A., Charron, C. S., Khan, N. A., et al. Fecal Metagenomics to Identify Biomarkers of Food Intake in Healthy Adults: Findings from Randomized, Controlled, Nutrition Trials. The Journal of Nutrition 154, 271–283 (2024).

59. Kingma, D. P. & Ba, J. Adam: A Method for Stochastic Optimization. Preprint at 10.48550/arXiv.1412.6980 (2017).

60. Wang, T. Predicting metabolite response to dietary intervention using deep learning. wt1005203/McMLP: v1.0.1. Zenodo 10.5281/zenodo.11113978 (2024).

