## Supplemental Information for "Predicting metabolite response to dietary intervention using deep learning"

### Supplementary Note

#### Existing machine learning methods

*Gradient-Boosting Regressor (GBR)*. GBR is a computational method used in a previous study that attempted to predict glycemic responses after food intake[3]. GBR uses a one-step approach and does not leverage the endpoint microbial compositions during method training.

- Data processing: The CLR (Centered Log-Ratio) transformation is applied to microbial abundances and the log10 transformation is applied to metabolite concentrations. Hyperparameter selection: Three hyperparameters are selected based on the 5-fold cross-validation results (the mean Spearman's rank correlation coefficients) on the training set: (1) the number of boosting iterations $N_{\mathrm{b}}$, (2) maximum depth $d$, and (3) learning rate $l$. $N_{\mathrm{b}}$ is selected from [1, 3, 10, 30, 100, 300, 1000]. $d$ is selected from [2, 4, 8, 10, 12]. $l$ is selected from [$10^{-4}$, $10^{-3}$, $10^{-2}$, $10^{-1}$, 1].

*Random Forest (RF)*. RF is a computational method used in a previous study that attempted to predict postprandial responses of blood triglyceride, glucose, and insulin after food intake[4]. RF uses a one-step approach and does not leverage the endpoint microbial compositions during method training.

- Data processing: The CLR (Centered Log-Ratio) transformation is applied to microbial abundances and the log10 transformation is applied to metabolite concentrations.
- Hyperparameter selection: Four hyperparameters are selected based on the 5-fold cross-validation results (the mean Spearman's rank correlation coefficients) on the training set: (1) the number of decision tree $N_{\mathrm{num\_of\_trees}}$, (2) maximum number of features $N_{\mathrm{features}}$, (3) minimum number of samples required to split an internal node $N_{\mathrm{min\ \_samples\_split}}$, and (4) minimum number of samples required to be at a leaf node $N_{\mathrm{min\ \_samples\_leaf}}$. $N_{\mathrm{num\_of\_trees}}$ is selected from [1, 10, 30, 100, 300, 1000]. $N_{\mathrm{features}}$ is selected from either "sqrt" (i.e., square root of number of features as the maximal number of features) or "log2" (i.e., log2 of number of features as the maximal number of features). $N_{\mathrm{min\ \_samples\_split}}$ is selected from [2, 5, 10, 15, 20]. $N_{\mathrm{min\ \_samples\_leaf}}$ is selected from [1, 2, 4, 6, 8].

**The difference between McMLP and previous methods (which predict the metabolomic profile based on the microbial composition at the same time).** There are several existing methods for the inference of metabolomic profiles from microbial compositions measured at the same time[35–38]. It has been shown that mNODE[38] displays the best performance among them. But they all require the microbial composition and metabolomic profile collected at the same time. In addition, their performances are good when the number of metabolites is large, especially for untargeted metabolomics. This is because most of the methods rely on techniques to reduce model complexity (e.g., regularization, the encoder-decoder architecture). However, for many dietary intervention datasets, only a few metabolite biomarkers were selected. In this context, we developed McMLP based on the overparametrized MLP ($N_{\mathrm{l}} = 6$ and $N_{\mathrm{h}} = 2048$) because the overparametrized deep-learning model (i.e., one with an extremely large number of model parameters) can generate better and more consistent performance than models with less capacity and more carefully tuned hyperparameters[44]. The overparameterized machine learning methods could have a better performance because their high capacity (i.e., more model parameters) can make them even simpler due to smoother function approximation and thus less likely to overfit[44].

**Synthetic data generated by customized MiCRM with periodical supplies of food resources and dilution.** Similar to the MiCRM[45], We customized the model to simulate the process of periodic supplies of food resources, consumption of food resources, growth of microbes, production and consumption of metabolites, as well as periodic dilution of the community. We used 20 microbial species, 20 food resources, and 20 metabolites in total in the pool. We separated food resources from metabolites, assuming that food resources can only be consumed while metabolites can be either consumed or produced. Prior to dietary intervention, one food resource (referred to as “food resource #1”) was not introduced, while the remaining 19 food resources were supplied. We then simulated the population dynamics and collected microbial relative abundances for surviving species and metabolite concentrations as baseline data for microbial composition and metabolomic profiles. Dietary intervention was simulated by adding food resource #1 at specific doses to communities composed of surviving species before the dietary intervention and calculating the new ecological steady state. After simulating the dietary intervention, we recorded microbial relative abundances and metabolite concentrations as the endpoint data. The concentration of food resource #1 supplied during the dietary intervention was used as an input variable in McMLP to represent the dietary intervention strategy. The same set of model parameters such as the consumption, production, and dilution rates is assumed when different samples in synthetic data were generated. Specially, the process for generating each sample can be divided into 3 steps:

(1) Sampling of an initial population before the dietary intervention: a fraction of microbial species are sampled from the metapopulation with a probability of $p_s(= 50\%)$ being selected to be present for each species. Similarly, a fraction of food resources except for food resource 1 that is introduced later are sampled from the metapopulation with a probability of $p_f(= 60\%)$ being selected to be supplied for each food resource.

(2) Simulation of population dynamics before the dietary intervention: after the present species and food resources are determined, we introduced fixed and predetermined amounts of food resources and simulated the population dynamics in the next 24 hours. 24 hours later, we diluted the community (microbial abundances, concentrations of food resources, and metabolite concentrations) by a constant factor of 10 and then refilled the community with the same predetermined amounts of food resources to begin another round of simulation. We call every 24 hours of dilution, feeding, and population growth a "feed-dilution cycle". We collected the microbial relative abundances and metabolite concentrations after 10 feed-dilution cycles of simulations as the baseline microbial composition and metabolomic profiles.

(3) Simulation of population dynamics after the dietary intervention: now we added food resource 1 (which is the intervened food resource during the diet intervention) to the previous food profile and reduced concentrations of other food resources proportionally to conserve the total concentration of supplied food resources, which form the new diet (i.e., the new combination of supplied food resources). We took the survived species and their abundances from the previous step and simulated the population dynamics in the next 10 feed-dilution cycles. After that, we recorded the microbial relative abundances and metabolite concentrations in simulations as the endpoint microbial composition and metabolomic profiles. The amount of introduced food resource 1 is also documented as the input variable for the dietary intervention strategy. An alternative version with 5 added food resources is also tried by introducing 5 food resources with the fixed total dose of 3 (i.e., the doses of all 5 food resources add up to 3, though the dose of each food resource can vary).

This 3-step sample-generating procedure is repeated many times to create independent samples in the synthetic data. For this study, we generated 250 samples in total and split the synthetic data fifty times with 80/20 ratio to generate fifty train-test pairs that can be used to reflect the variation in predictive performance. During each feed-dilution cycle, microbial species $i$ consumes all food resources and metabolites it is capable of. Once microbial species $i$ consumes, all consumed food resources and metabolites contribute to two parts: (1) a fraction $1 - l$ of total consumption amount contributes to microbial growth, and (2) the remaining fraction $l$ of total consumption amount is converted to other metabolite byproducts, with their byproduct generation fluxes encoded by a pre-determined vector for species $i$ (denoted as $P_{i\beta}$). $\sum_\beta P_{i\beta} = 1$ is imposed to guarantee the conservation of total concentration of metabolites. The overall population dynamics for the concentration of food resource $R_\alpha$, abundance of microbial species $N_i$, and concentration of metabolite $M_\beta$ can be written as follows:

$$\frac{dR_\alpha}{dt} = -\sum_i a_{i\alpha} N_i R_\alpha,$$

$$\frac{dN_i}{dt} = \frac{(1-l)(\sum_\gamma a_{i\gamma} N_i R_\gamma + \sum_\delta b_{i\delta} N_i M_\delta)}{Y},$$

$$\frac{dM_\beta}{dt} = -\sum_i b_{i\beta} N_i M_\beta + l \sum_i P_{i\beta} (\sum_\gamma a_{i\gamma} N_i R_\gamma + \sum_\delta b_{i\delta} N_i M_\delta),$$

where $a_{i\alpha}$ is the consumption rate of food resource $\alpha$ by the species $i$, $b_{i\beta}$ is the consumption rate of metabolite $\beta$ by the species $i$, and $Y$ is the yield. The exact values for those model parameters and how they are assigned are as follows:

- $a_{i\alpha}$ is assumed to have a connectance of 50%, meaning that each element in $a_{i\alpha}$ is non-zero with a probability of 50%. For each non-zero element, its value is randomly drawn from the uniform distribution between 0 and 10. After the assignment of values, the consumption $a_{i\alpha}$ is divided by the number of food resources one species can consume. We imposed this division to avoid the dominance of generalist specialists that consume almost all food resources with a penalty. $b_{i\beta}$ is assigned in the same way as $a_{i\alpha}$.
- $P_{i\beta}$ is assumed to have a connectance of 50%. For each non-zero element, its value is randomly drawn from the uniform distribution between 0 and 1. If $b_{i\beta} \neq 0$ for a pair of metabolite $\alpha$ and species $i$, we set $P_{i\beta} = 0$ for a pair of metabolite $\alpha$ and species $i$. We impose this condition to avoid one metabolite being consumed and produced by the same species at the same time. Then $\sum_\beta P_{i\beta} = 1$ is imposed.
- The byproduct fraction $l = 0.5$ for all scenarios.
- The yield $Y = 1$ for simplicity.
- The dilution factor is 10.

### Supplementary Figures

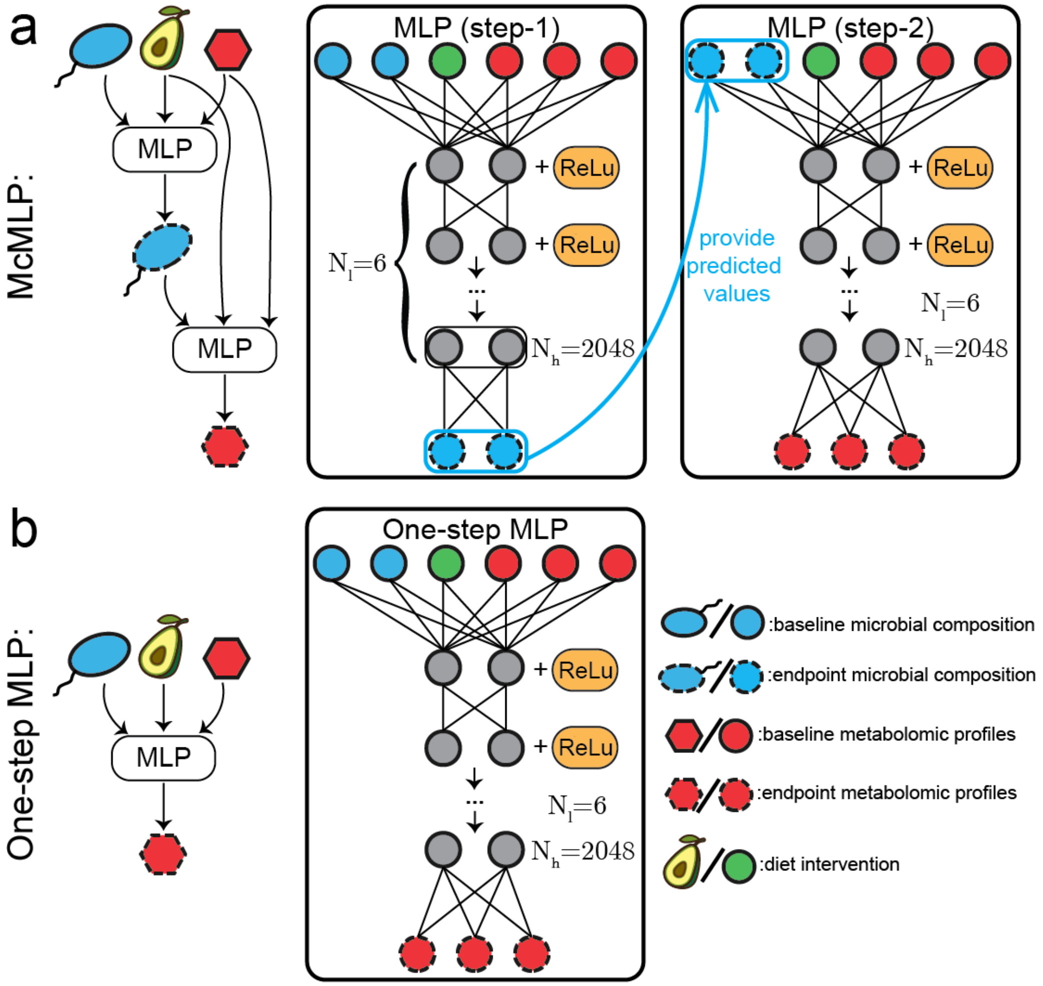


Supplementary Figure 1: **The architecture of McMLP and one-step MLP.** To predict endpoint metabolomic profiles (i.e., metabolomic profiles after the dietary interventions) based on the baseline microbial compositions (i.e., microbial compositions before the dietary intervention), dietary intervention strategy, and baseline metabolomic profiles, we can have two strategies: McMLP and one-step MLP. Across panels, microbial species and their relative abundances are colored blue, dietary resources and their intervention doses are colored green, and metabolites and their concentrations are colored red. Icons associated with baseline/endpoint data are bounded by solid black/dashed lines respectively. The number of hidden layers is $N_l$ and the number of nodes in each hidden layer is $N_h$. **a,** The model architecture of McMLP comprises two coupled MLPs. The first MLP (step 1) predicts the endpoint microbial compositions based on the baseline data and the dietary intervention strategy. Then the predicted endpoint microbial compositions from the first MLP are provided as the input variables for the second MLP (step 2). The second MLP combines the predicted endpoint microbial compositions, dietary intervention strategy, and baseline metabolomic profiles to finally predict the endpoint metabolomic profiles. **b,** By contrast, the one-step MLP directly predicts endpoint metabolomic profiles based on the baseline data and the dietary intervention strategy.

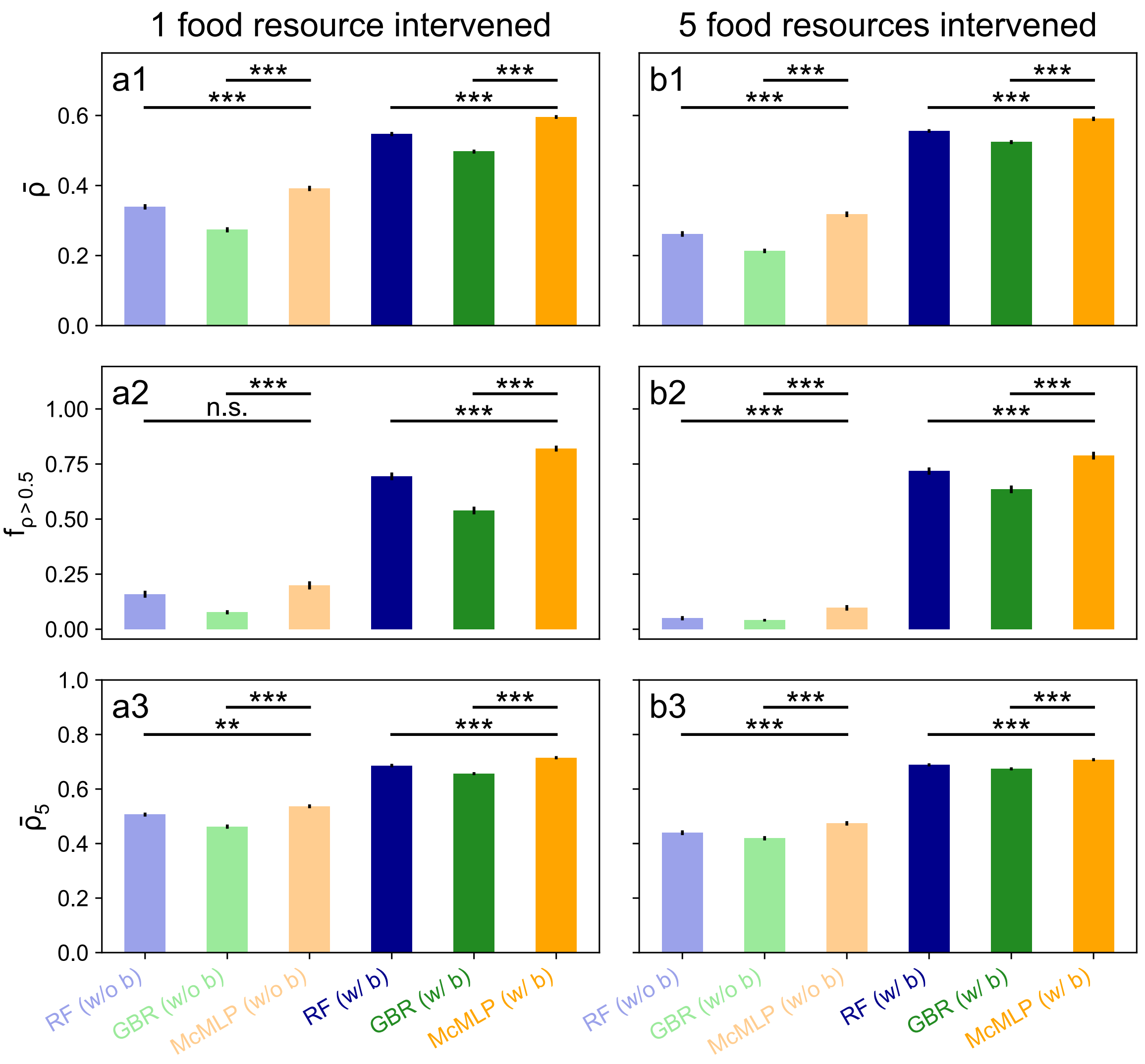


Supplementary Figure 2: **The predictive performance of McMLP is superior to other methods when more food resources are introduced during the dietary intervention.** Synthetic data are generated from microbial consumer-resource models. For each method, we either included (i.e., the case of "w/ b") or did not (i.e., the case of "w/o b") include baseline metabolomic profiles as input variables. Each method with a particular input data combination is colored in the same way across all panels. Standard errors are computed based on 50 train-test splits and shown in all panels (as solid black vertical lines or transparent areas around their means). To compare different methods, we adopted three metrics: the mean Spearman Correlation Coefficient (SCC) $\bar{\rho}$, the fraction of metabolites with SCCs greater than 0.5 (denoted as $f_{\rho>0.5}$), and the mean SCC of the top-5 predicted metabolites $\bar{\rho}_5$. Error bars denote the standard error (n=50). **a1-a3,** Comparison of predictive performance when 1 food resource is introduced during the dietary intervention, the intervention dose is 3, and the training sample size equals 50. **b1-b3,** Comparison of predictive performance when 5 food resources are introduced during the dietary intervention, the intervention dose is 3, and the training sample size equals 50. All statistical analyses were performed using the two-sided Wilcoxon signed-rank test. P values obtained from the test are divided into four groups: (1) $p > 0.05$ (n.s.), (2) $0.01 < p \leq 0.05$ (*), (3) $10^{-3} < p \leq 0.01$ (**), and (4) $10^{-4} < p \leq 10^{-3}$ (***).

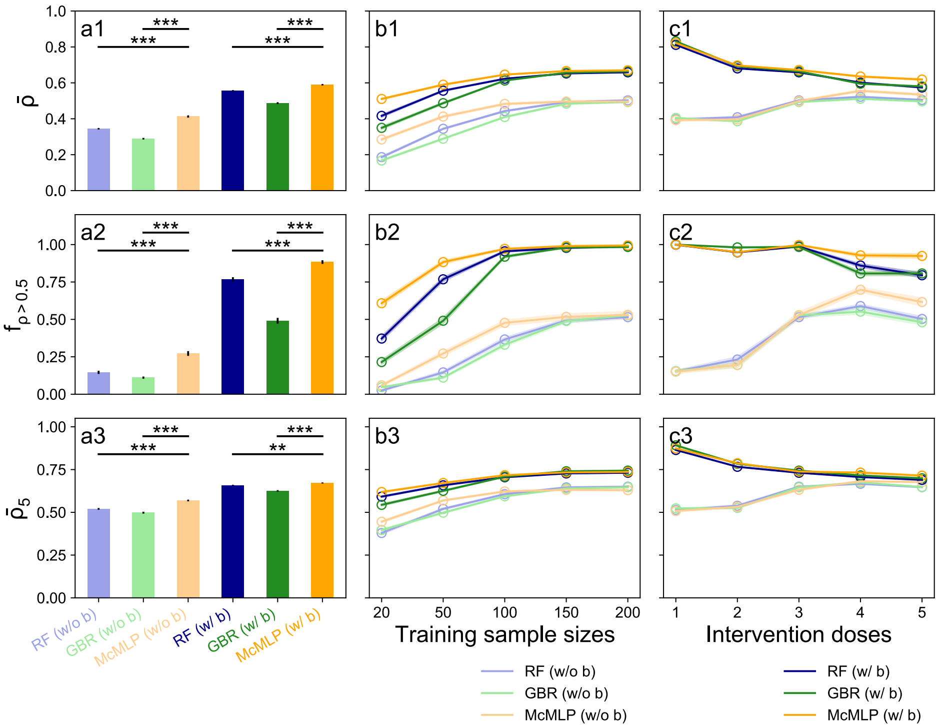


Supplementary Figure 3: **McMLP provides better predictive power than previously developed computational methods on synthetic data generated by MiCRM when training data across repeats do not overlap.** In these simulations, MiCRM is used to freshly generate all training data for each training repeat, ensuring that there is no overlap of training data between different repeats. However, the same set of test data is employed across all repeats to assess the predictive performance of each repeat. Three computational methods are compared: Random Forest (RF), Gradient Boosting Regressor (GBR), and McMLP. For each method, we either included ("w/ b" label) or did not include ("w/o b" label) baseline metabolomic profiles as input variables. Each method with a particular combination of input data is colored the same way in all panels. Standard errors are computed based on fifty training repeats and shown in all panels (as solid black vertical lines or transparent areas around their means). To compare different methods, we adopted three metrics: the mean Spearman Correlation Coefficient (SCC) $\bar{\rho}$, the fraction of metabolites with SCCs greater than 0.5 (denoted as $f_{\rho>0.5}$), and the mean SCC of the top-5 predicted metabolites $\bar{\rho}_5$. Error bars denote the standard error (n=50). **a1-a3,** For the synthetic data with intervention dose of 3 and 50 training samples, McMLP provides the best performance for all three metrics regardless of whether the baseline metabolomic profiles are included or not. **b1-b3,** When the intervention dose is 3, the predictive performance of all methods gets better and closer to each other as the training sample size increases. Including baseline metabolomic profiles also helps to improve the prediction. **c1-c3,** When 200 training samples are used, the performance gap between including and not including baseline metabolomic profiles shrinks as the intervention dose increases. All statistical analyses were performed using the two-sided Wilcoxon signed-rank test. P values obtained from the test are divided into four groups: (1) $p > 0.05$ (n.s.), (2) $0.01 < p \leq 0.05$ (*), (3) $10^{-3} < p \leq 0.01$ (**), and (4) $10^{-4} < p \leq 10^{-3}$ (***).

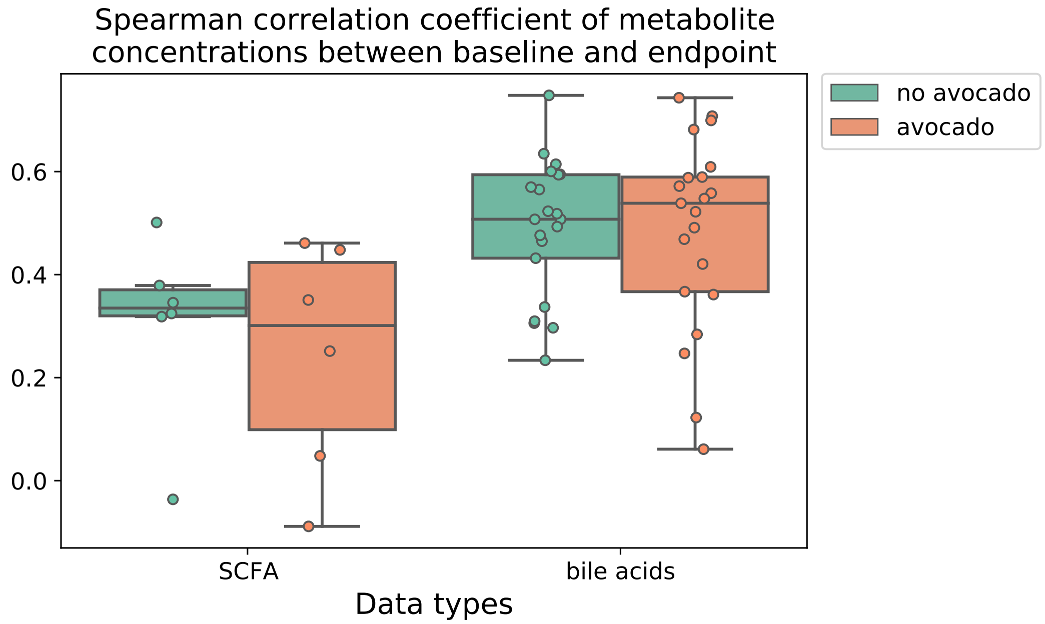


Supplementary Figure 4: **The mean Spearman Correlation coefficients of bile acids between baseline and endpoint is larger than that of SCFAs in the data from an avocado intervention study[28].** Error bars denote the standard error (n=5). In all boxplots, the middle black line is the median, the box extends from the first quartile (Q1) to the third quartile (Q3) of the data, the black whiskers extend from the box by 1.5 × IQR (where IQR is the interquartile range), and outlier unfilled black circles are those beyond the range defined by two whiskers.

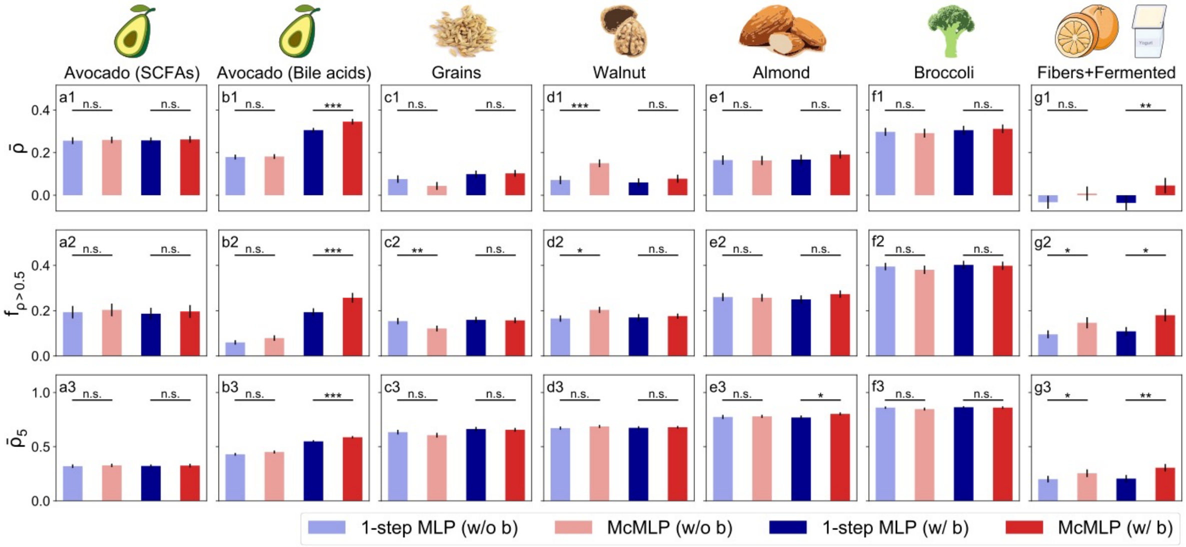


Supplementary Figure 5: **McMLP is better than one-step MLP in terms of predicting endpoint metabolomic profiles on real data from dietary intervention studies.** The 1-step MLP does not predict endpoint microbial compositions first. Instead, it directly used the baseline microbial compositions, baseline metabolomic profiles, and dietary intervention strategy to predict the endpoint metabolomic profiles. The one-step McMLP uses the same number of layers and nodes as one step in McMLP ($N_{\mathrm{l}} = 6$ and $N_{\mathrm{h}} = 2048$). For both methods, we either included (i.e., the case of "w/ b") or did not (i.e., the case of "w/o b") include baseline metabolomic profiles as input variables. Each method with a particular input data combination is colored in the same way across all panels. Standard errors are computed based on 50 train-test splits and shown in all panels (solid black vertical lines). To compare different methods, we adopted three metrics: the mean Spearman Correlation Coefficient (SCC) $\bar{\rho}$, the fraction of metabolites with SCCs greater than 0.5 (denoted as $f_{\rho>0.5}$), and the mean SCC of the top-5 predicted metabolites $\bar{\rho}_5$. Error bars denote the standard error (n=50). **a1-a3,** Comparison of performance of predicting SCFAs in the data from an avocado intervention study[28]. **b1-b3,** Comparison of performance of predicting bile acids in the data from an avocado intervention study[28]. **c1-c3,** Comparison of predictive performance on the data from a grain intervention study[39]. **d1-d3,** Comparison of predictive performance on the data from a walnut intervention study[27]. **e1-e3,** Comparison of predictive performance on the data from an almond intervention study[40]. **f1-f3,** Comparison of predictive performance on the data from a broccoli intervention study[41]. **g1-g3,** Comparison of predictive performance on the data from a high-fiber food or fermented food intervention study[34]. All statistical analyses were performed using the two-sided Wilcoxon signed-rank test. P values obtained from the test are divided into four groups: (1) $p > 0.05$ (n.s.), (2) $0.01 < p \leq 0.05$ (*), (3) $10^{-3} < p \leq 0.01$ (**), and (4) $10^{-4} < p \leq 10^{-3}$ (***).

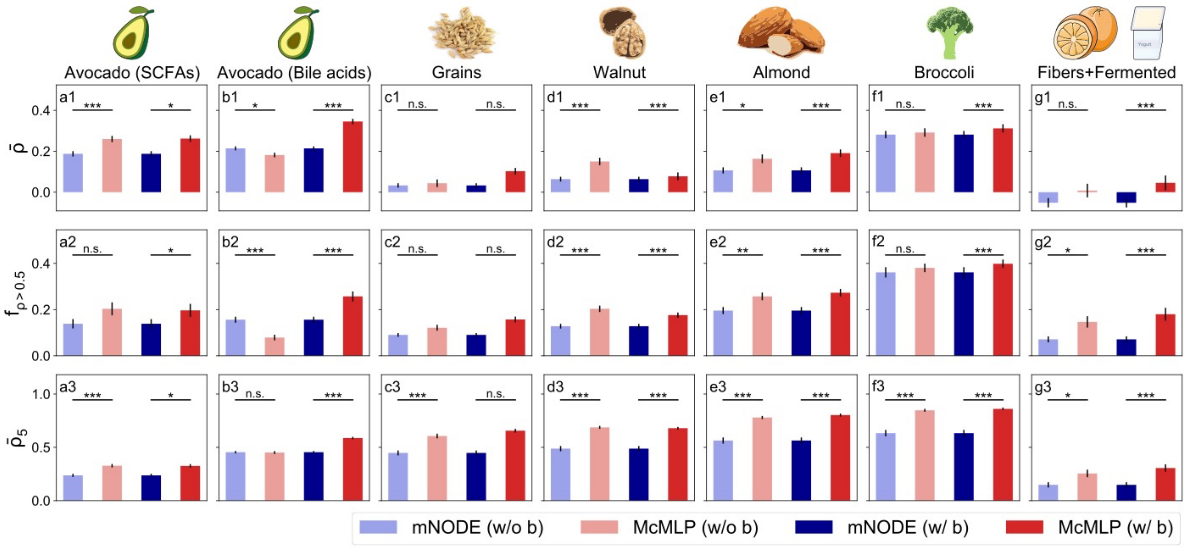


Supplementary Figure 6: **McMLP is superior to mNODE on real data from dietary intervention studies.** mNODE has been shown to be the state-of-the-art method for predicting metabolomic profiles based on the microbial compositions measured at the same time[38]. For both methods, we either included (i.e., the case of "w/ b") or did not (i.e., the case of "w/o b") include baseline metabolomic profiles as input variables. Each method with a particular input data combination is colored in the same way across all panels. Standard errors are computed based on 50 train-test splits and shown in all panels (solid black vertical lines). To compare different methods, we adopted three metrics: the mean Spearman Correlation Coefficient (SCC) $\bar{\rho}$, the fraction of metabolites with SCCs greater than 0.5 (denoted as $f_{\rho>0.5}$), and the mean SCC of the top-5 predicted metabolites $\bar{\rho}_5$. Error bars denote the standard error (n=50). **a1-a3,** Comparison of performance of predicting SCFAs in the data from an avocado intervention study[28]. **b1-b3,** Comparison of performance of predicting bile acids in the data from an avocado intervention study[28]. **c1-c3,** Comparison of predictive performance on the data from a grain intervention study[39]. **d1-d3,** Comparison of predictive performance on the data from a walnut intervention study[27]. **e1-e3,** Comparison of predictive performance on the data from an almond intervention study[40]. **f1-f3,** Comparison of predictive performance on the data from a broccoli intervention study[41]. **g1-g3,** Comparison of predictive performance on the data from a high-fiber food or fermented food intervention study[34]. All statistical analyses were performed using the two-sided Wilcoxon signed-rank test. P values obtained from the test are divided into four groups: (1) $p > 0.05$ (n.s.), (2) $0.01 < p \leq 0.05$ (*), (3) $10^{-3} < p \leq 0.01$ (**), and (4) $10^{-4} < p \leq 10^{-3}$ (***).

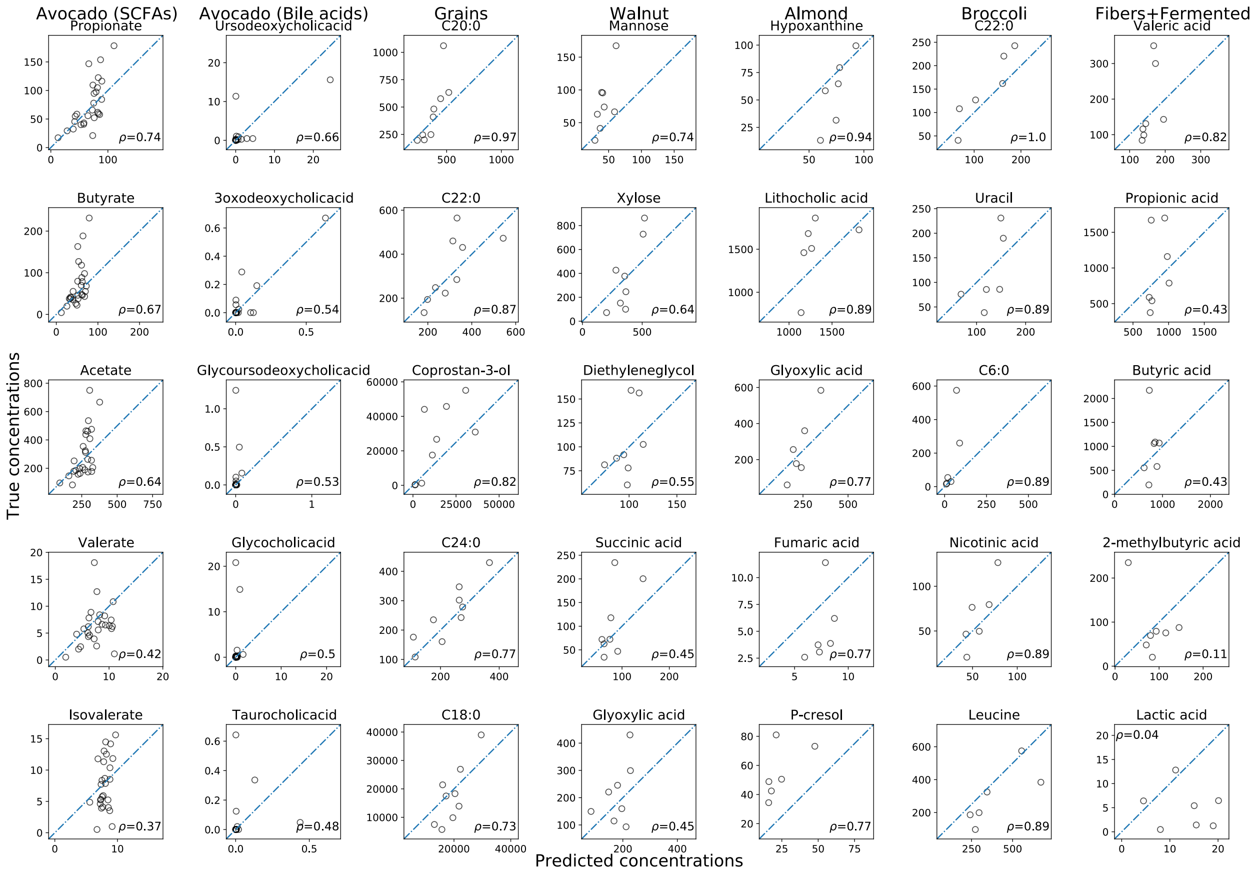


Supplementary Figure 7: **Comparison between predicted and true metabolite concentrations on the test set of the real data from six dietary intervention studies.** For each dataset, the top-5 predicted metabolites (i.e., the predicted metabolites with the highest Spearman Correlation Coefficients $\rho$) are shown from top to bottom.

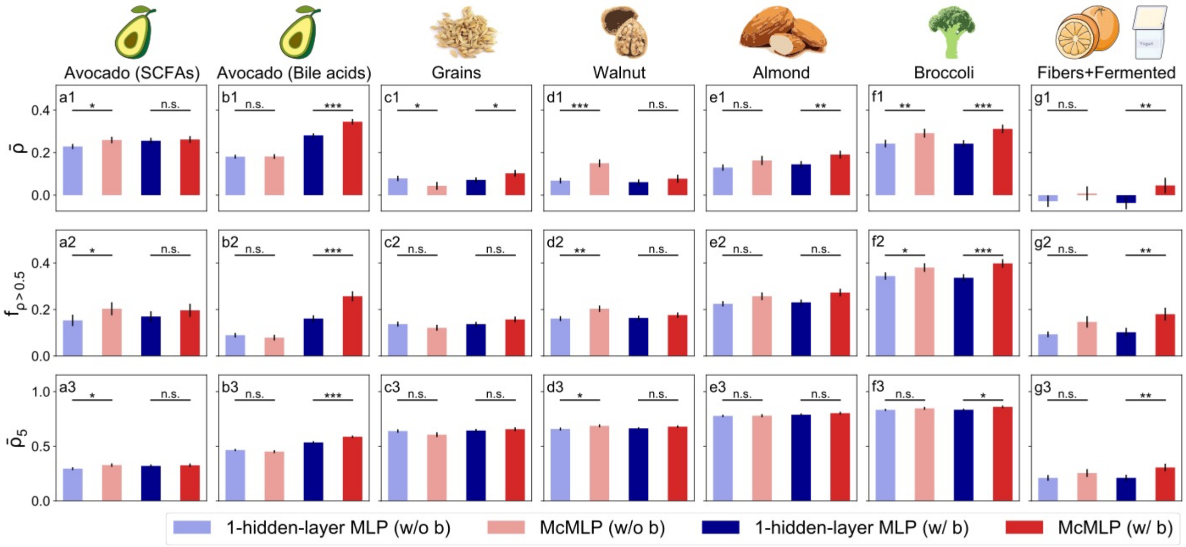


Supplementary Figure 8: **McMLP is superior to 1-hidden-layer MLP in terms of predicting endpoint metabolomic profiles on real data from dietary intervention studies.** The 1-hidden-layer MLP only has one hidden layer, while other parameters, hyperparameters, and the training protocol are the same as that of McMLP. For both methods, we either included (i.e., the case of "w/ b") or did not (i.e., the case of "w/o b") include baseline metabolomic profiles as input variables. Each method with a particular input data combination is colored in the same way across all panels. Standard errors are computed based on 50 train-test splits and shown in all panels (solid black vertical lines). To compare different methods, we adopted three metrics: the mean Spearman Correlation Coefficient (SCC) $\bar{\rho}$, the fraction of metabolites with SCCs greater than 0.5 (denoted as $f_{\rho>0.5}$), and the mean SCC of the top-5 predicted metabolites $\bar{\rho}_5$ . Error bars denote the standard error (n=50). **a1-a3,** Comparison of performance of predicting SCFAs in the data from an avocado intervention study[28]. **b1-b3,** Comparison of performance of predicting bile acids in the data from an avocado intervention study[28]. **c1-c3,** Comparison of predictive performance on the data from a grain intervention study[39]. **d1-d3,** Comparison of predictive performance on the data from a walnut intervention study[27]. **e1-e3,** Comparison of predictive performance on the data from an almond intervention study[40]. **f1-f3,** Comparison of predictive performance on the data from a broccoli intervention study[41]. **g1-g3,** Comparison of predictive performance on the data from a high-fiber food or fermented food intervention study[34]. All statistical analyses were performed using the two-sided Wilcoxon signed-rank test. P values obtained from the test are divided into four groups: (1) $p > 0.05$ (n.s.), (2) $0.01 < p \leq 0.05$ (*), (3) $10^{-3} < p \leq 0.01$ (**), and (4) $10^{-4} < p \leq 10^{-3}$ (***).

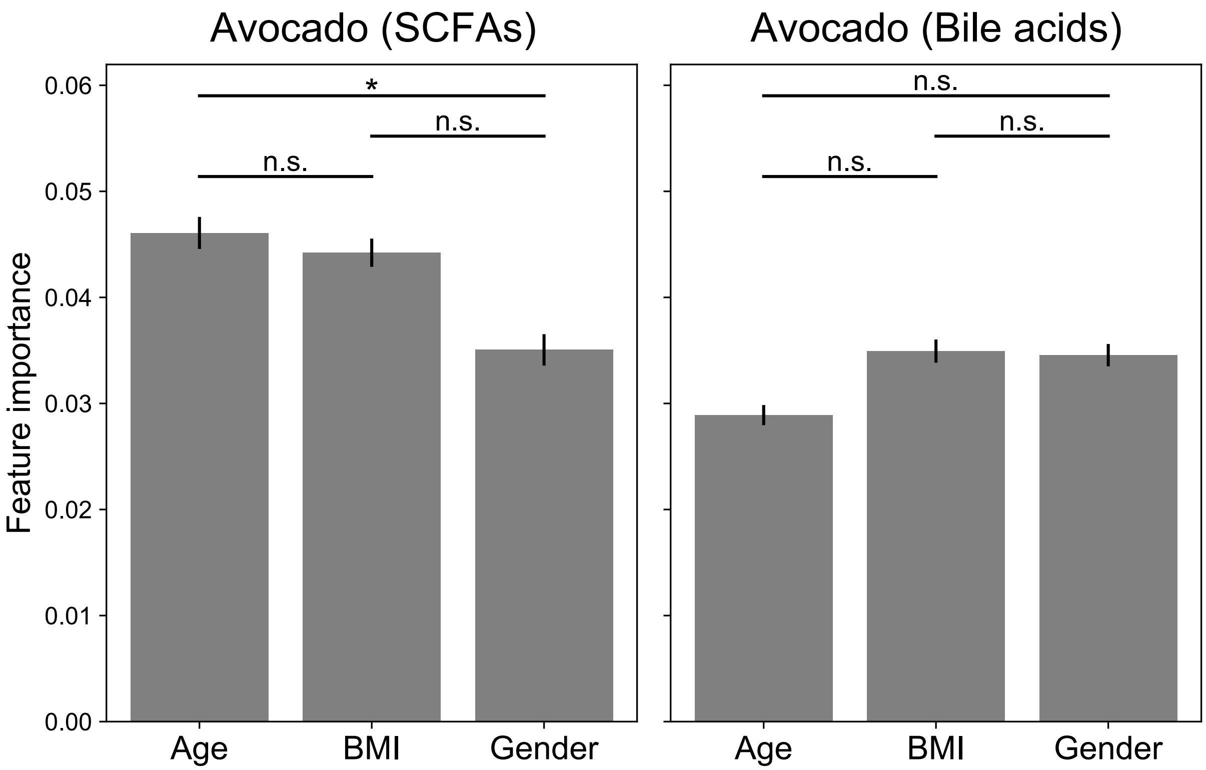


Supplementary Figure 9: **The feature importance of the covariates (age, BMI, and gender) included in the input of McMLP.** The feature importance is measured by the reduction in the mean Spearman Correlation Coefficient (SCC) $\bar{\rho}$ when a covariate is set as its mean value in the input. All results are derived from McMLP. Standard errors are computed based on fifty random train-test splits and shown in all panels (solid black vertical lines). Error bars denote the standard error (n=50). **a1-a3,** Comparison of the performance in predicting SCFAs on the data from the avocado intervention study[28]. **b1-b3,** Comparison of performance in predicting bile acids on the data from the avocado intervention study[28]. All statistical analyses were performed using the two-sided Wilcoxon signed-rank test. P values obtained from the test are divided into four groups: (1) $p > 0.05$ (n.s.), (2) $0.01 < p \leq 0.05$ (*), (3) $10^{-3} < p \leq 0.01$ (**), and (4) $10^{-4} < p \leq 10^{-3}$ (***).

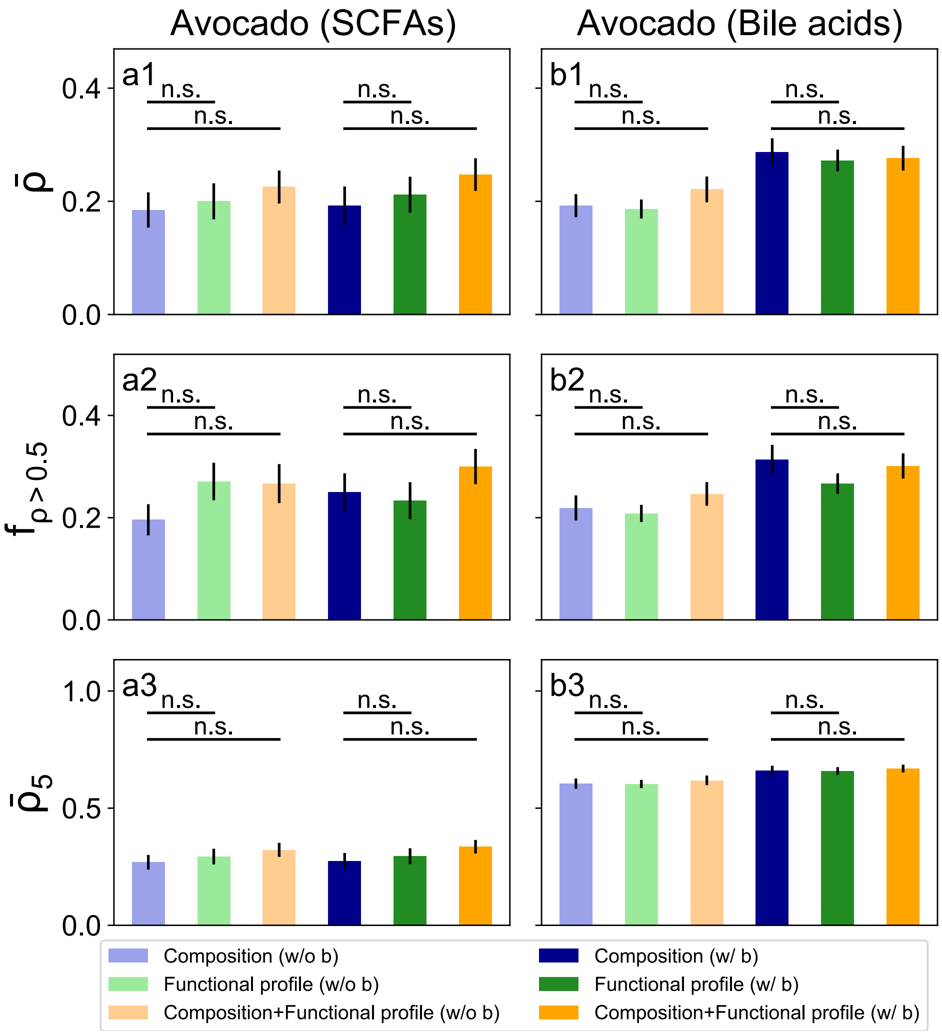


Supplementary Figure 10: **Using the microbial compositions from the 16S sequencing and/or using the functional profiles from the shotgun sequencing in the input of McMLP achieves similar predictive performance in terms of predicting endpoint metabolomic profiles on real data from the avocado intervention study.** All results are derived from McMLP. We either included ("w/ b" label) or did not include ("w/o b" label) baseline metabolomic profiles as input variables. Each method with a particular combination of input data is colored the same in all panels. Standard errors are computed based on fifty random train-test splits and shown in all panels (solid black vertical lines). To compare different methods, we adopted three metrics: the mean Spearman Correlation Coefficient (SCC) $\bar{\rho}$, the fraction of metabolites with SCCs greater than 0.5 (denoted as $f_{\rho>0.5}$), and the mean SCC of the top-5 predicted metabolites $\bar{\rho}_5$. Error bars denote the standard error (n=50). **a1-a3,** Comparison of the performance in predicting SCFAs on the data from the avocado intervention study[28]. **b1-b3,** Comparison of performance in predicting bile acids on the data from the avocado intervention study[28]. All statistical analyses were performed using the two-sided Wilcoxon signed-rank test. P values obtained from the test are divided into four groups: (1) $p > 0.05$ (n.s.), (2) $0.01 < p \leq 0.05$ (*), (3) $10^{-3} < p \leq 0.01$ (**), and (4) $10^{-4} < p \leq 10^{-3}$ (***).

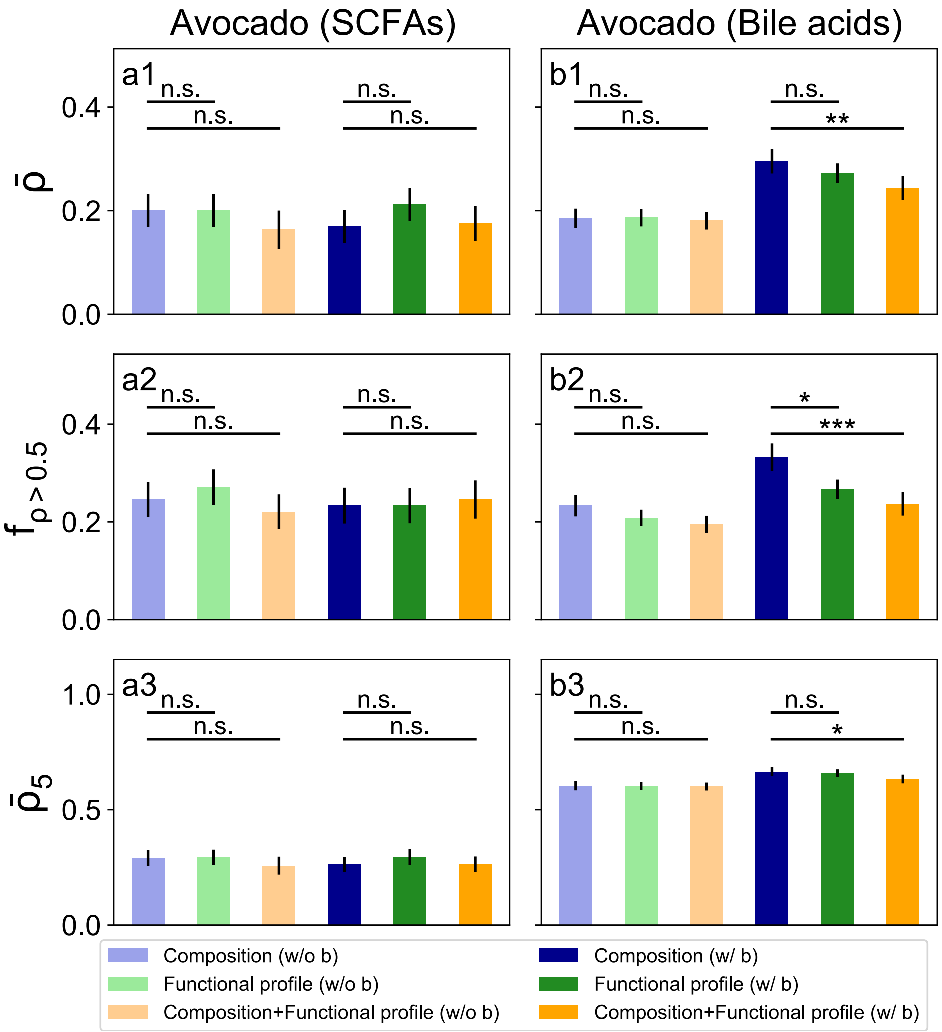


Supplementary Figure 11: **Using the microbial compositions from the shotgun sequencing and/or using the functional profiles from the shotgun sequencing in the input of McMLP achieves similar predictive performance in terms of predicting endpoint metabolomic profiles on real data from the avocado intervention study.** All results are derived from McMLP. We either included ("w/ b" label) or did not include ("w/o b" label) baseline metabolomic profiles as input variables. Each method with a particular combination of input data is colored the same in all panels. Standard errors are computed based on fifty random train-test splits and shown in all panels (solid black vertical lines). To compare different methods, we adopted three metrics: the mean Spearman Correlation Coefficient (SCC) $\bar{\rho}$, the fraction of metabolites with SCCs greater than 0.5 (denoted as $f_{\rho>0.5}$), and the mean SCC of the top-5 predicted metabolites $\bar{\rho}_5$. Error bars denote the standard error (n=50). **a1-a3,** Comparison of the performance in predicting SCFAs on the data from the avocado intervention study[28]. **b1-b3,** Comparison of performance in predicting bile acids on the data from the avocado intervention study[28]. All statistical analyses were performed using the two-sided Wilcoxon signed-rank test. P values obtained from the test are divided into four groups: (1) $p > 0.05$ (n.s.), (2) $0.01 < p \le 0.05$ (*), (3) $10^{-3} < p \le 0.01$ (**), and (4) $10^{-4} < p \le 10^{-3}$ (***).

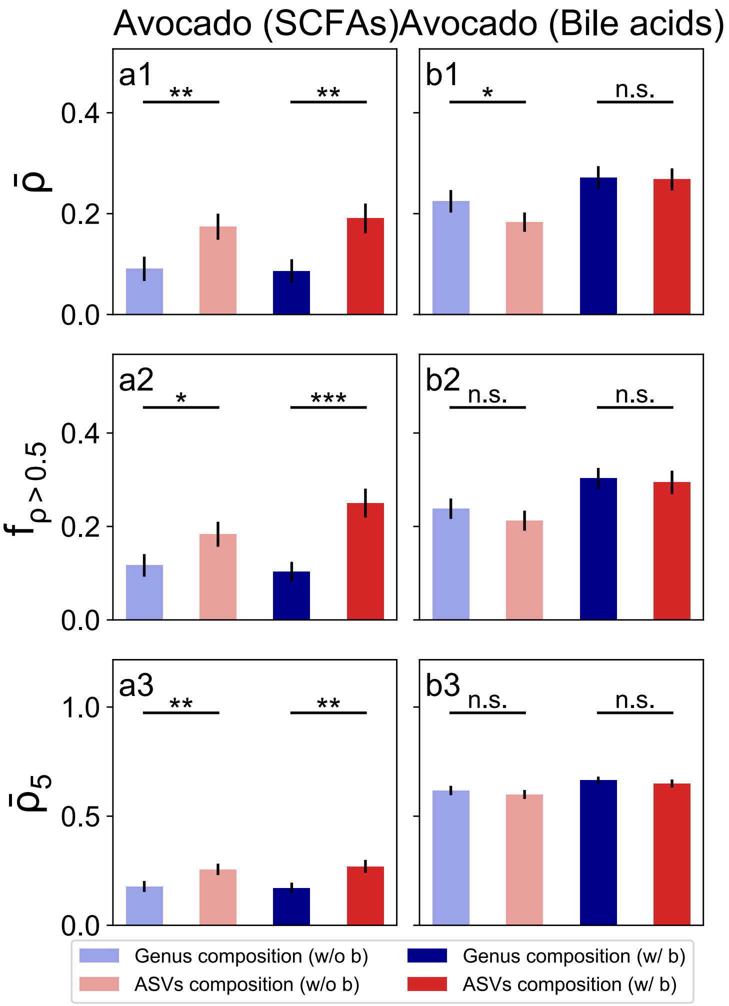


Supplementary Figure 12: **Using the ASV-level microbial compositions from the 16S sequencing in the input of McMLP achieves better predictive performance than using the genus-level microbial compositions in terms of predicting endpoint metabolomic profiles on real data from the avocado intervention study.** All results are derived from McMLP. We either included ("w/ b" label) or did not include ("w/o b" label) baseline metabolomic profiles as input variables. Each method with a particular combination of input data is colored the same in all panels. Standard errors are computed based on fifty random train-test splits and shown in all panels (solid black vertical lines). To compare different methods, we adopted three metrics: the mean Spearman Correlation Coefficient (SCC) $\bar{\rho}$, the fraction of metabolites with SCCs greater than 0.5 (denoted as $f_{\rho>0.5}$), and the mean SCC of the top-5 predicted metabolites $\bar{\rho}_5$. Error bars denote the standard error (n=50). **a1-a3,** Comparison of the performance in predicting SCFAs on the data from the avocado intervention study[28]. **b1-b3,** Comparison of performance in predicting bile acids on the data from the avocado intervention study[28]. All statistical analyses were performed using the two-sided Wilcoxon signed-rank test. P values obtained from the test are divided into four groups: (1) $p > 0.05$ (n.s.), (2) $0.01 < p \leq 0.05$ (*), (3) $10^{-3} < p \leq 0.01$ (**), and (4) $10^{-4} < p \leq 10^{-3}$ (***).

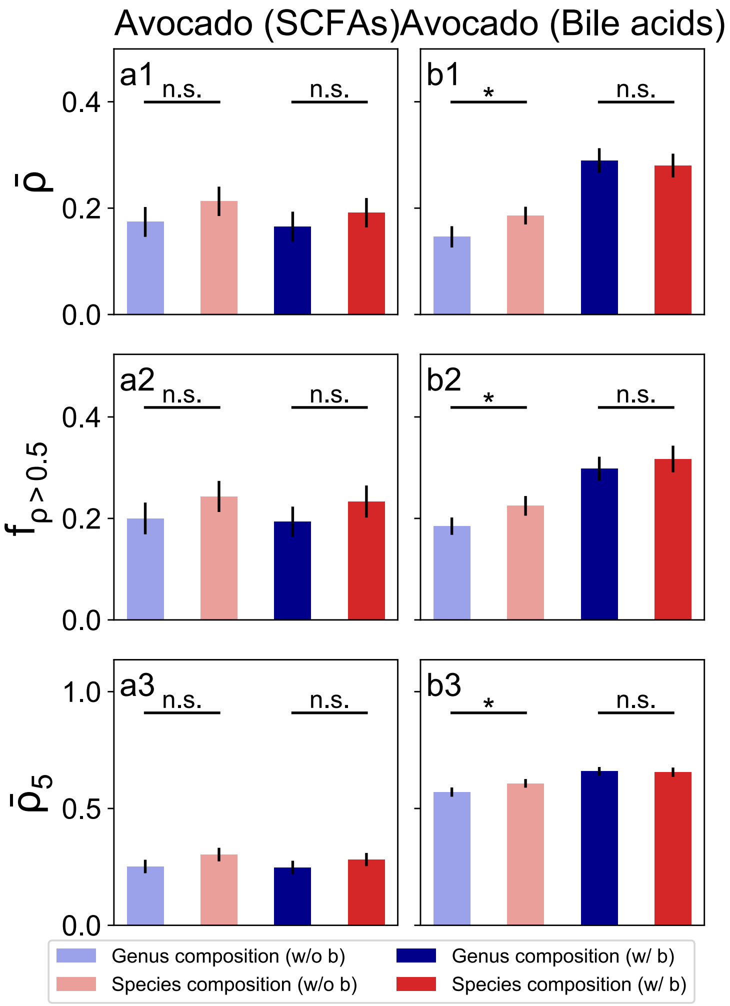


Supplementary Figure 13: **Using the species-level microbial compositions from the shotgun sequencing in the input of McMLP achieves better predictive performance than using the genus-level microbial compositions in terms of predicting endpoint metabolomic profiles on real data from the avocado intervention study.** All results are derived from McMLP. We either included ("w/ b" label) or did not include ("w/o b" label) baseline metabolomic profiles as input variables. Each method with a particular combination of input data is colored the same in all panels. Standard errors are computed based on fifty random train-test splits and shown in all panels (solid black vertical lines). To compare different methods, we adopted three metrics: the mean Spearman Correlation Coefficient (SCC) $\bar{\rho}$, the fraction of metabolites with SCCs greater than 0.5 (denoted as $f_{\rho>0.5}$), and the mean SCC of the top-5 predicted metabolites $\bar{\rho}_5$. Error bars denote the standard error (n=50). **a1-a3,** Comparison of the performance in predicting SCFAs on the data from the avocado intervention study[28]. **b1-b3,** Comparison of performance in predicting bile acids on the data from the avocado intervention study[28]. All statistical analyses were performed using the two-sided Wilcoxon signed-rank test. P values obtained from the test are divided into four groups: (1) $p > 0.05$ (n.s.), (2) $0.01 < p \leq 0.05$ (*), (3) $10^{-3} < p \leq 0.01$ (**), and (4) $10^{-4} < p \leq 10^{-3}$ (***).

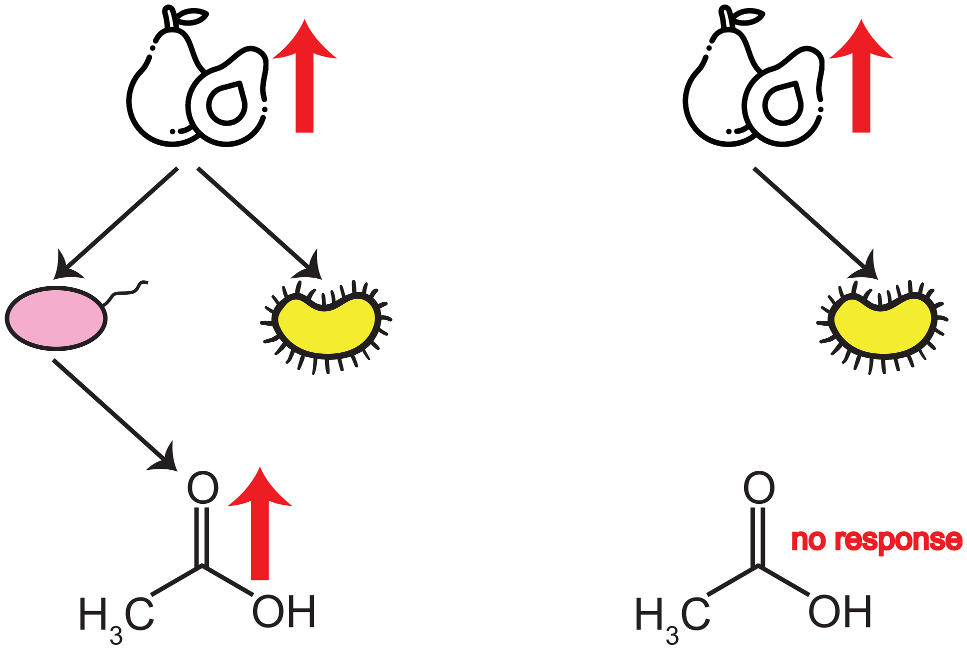


Supplementary Figure 14: **Whether the diet intervention can induce correct metabolite responses depends on species that can both consume the constituents of introduced food resources and produce the health-beneficial metabolite.**

**Supplementary Data Legends**

**Supplementary Data 1:** A table of sensitivity values for the avocado study[28].

**Supplementary Data 2:** A table of sensitivity values for the grain study[39].

**Supplementary Data 3:** A table of sensitivity values for the walnut study[27].

**Supplementary Data 4:** A table of sensitivity values for the almond study[40].

**Supplementary Data 5:** A table of sensitivity values for the broccoli study[41].

**Supplementary Data 6:** A table of sensitivity values for the study of high-fiber foods or fermented foods[34].
